# A genome-wide screen identifies SCAI as a modulator of the UV-induced replicative stress response in human cells

**DOI:** 10.1101/2022.01.17.476707

**Authors:** Jean-François Lemay, Edlie St-Hilaire, Sari Gezzar-Dandashi, Mary McQuaid, Daryl A. Ronato, Yuandi Gao, François Bélanger, Christina Sawchyn, Aimé Boris Kimenyi Ishimwe, Frédérick A. Mallette, Jean-Yves Masson, Elliot A. Drobetsky, Hugo Wurtele

## Abstract

Helix-destabilizing DNA lesions induced by environmental mutagens such as UV light cause genomic instability by strongly blocking the progression of DNA replication forks (RF). At blocked RF, single-stranded DNA (ssDNA) accumulates and is rapidly bound by Replication Protein A (RPA) complexes. Such stretches of RPA-ssDNA constitute platforms for recruitment/activation of critical factors that promote DNA synthesis restart. However, during periods of severe replicative stress, RPA availability may become limiting due to inordinate sequestration of this multifunctional complex on ssDNA, thereby negatively impacting multiple vital RPA-dependent processes. Here, we performed a genome-wide screen to identify factors which restrict the accumulation of RPA-ssDNA during UV-induced replicative stress. While this approach revealed some expected “hits” acting in pathways such as nucleotide excision repair, translesion DNA synthesis, and the intra-S phase checkpoint, it also identifed SCAI, whose role in the replicative stress response was previously unappreciated. Upon UV exposure, SCAI knock-down caused elevated accumulation of RPA-ssDNA during S phase, accompanied by reduced cell survival and compromised RF progression. These effects were independent of the previously reported role of SCAI in 53BP1-dependent DNA double-strand break repair. We also found that SCAI colocalized with stalled RF, and that its depletion promoted nascent DNA degradation. Finally, we (i) provide evidence that EXO1 is the major nuclease underlying ssDNA formation and consequent DNA replication defects in SCAI knockout cells and, consistent with this, (ii) demonstrate that SCAI inhibits EXO1 activity on a ssDNA gap *in vitro*. Taken together, our data establish SCAI as a novel regulator of the replicative stress response in human cells.

## INTRODUCTION

A variety of ubiquitous environmental genotoxins and chemotherapeutic drugs generate helix-destabilizing DNA adducts, e.g., solar UV-induced cyclobutane pyrimidine dimers (CPD) and 6-4 pyrimidine-pyrimidone photoproducts (6-4PP). If not efficiently removed by nucleotide excision repair (NER), such adducts block the progression of advancing replicative DNA polymerases. This, in turn, creates a state of “DNA replication stress” that precludes timely completion of S phase with potential genotoxic and carcinogenic consequences (Zeman and Cimprich, 2014). In order to alleviate these outcomes, i.e., to promote DNA synthesis restart, cells can enlist any among multiple DNA damage tolerance pathways to bypass replication-blocking lesions, including (i) error-free homologous recombination-dependent template switching (Branzei and Foiani, 2007), or (ii) error-prone translesion synthesis (TLS) following recruitment of specialized DNA polymerases to stalled replication forks (RF) (Goodman and Woodgate, 2013). In addition, Rad51-dependent replication fork reversal can promote reannealing of nascent DNA (Neelsen and Lopes, 2015; Zellweger et al., 2015). This brings replication-blocking lesions back into double-stranded DNA, thereby providing an opportunity to repair the lesion prior to eventual resumption of normal DNA replication. Recent evidence also demonstrates that repriming beyond damaged bases can also be used to allow continuation of DNA replication fork progression (Quinet et al., 2021).

Following genotoxin exposure, single-stranded DNA (ssDNA) generated at stalled RF is avidly bound by heterotrimeric Replication Protein A complexes (RPA) (Branzei and Foiani, 2009). This not only protects the ssDNA from degradation, but such RPA-bound ssDNA (hereafter RPA-ssDNA) also signals rapid activation of ATM and Rad3-related (ATR) kinase, the master regulator of intra S phase checkpoint signaling (Oakley and Patrick, 2010). ATR phosphorylates a multitude of substrates that cooperate to mitigate DNA replication stress by i) forestalling excessive accumulation of ssDNA at, and stabilizing, stalled RFs (Sogo et al., 2002; Zeman and Cimprich, 2014) and ii) preventing further RF blockage by repressing the activation of new origins of replication (Branzei and Foiani, 2009; Santocanale and Diffley, 1998). In addition, RPA is recruited to all active replication origins and advancing RF in the absence of genotoxic insult, where it coats/protects ssDNA resulting from normal MCM helicase activity (Diffley, 2004). In view of the above, maintaining an adequate supply of RPA during S phase, irrespective of whether or not cells are exposed to DNA damaging agents, is essential for timely completion of DNA synthesis (Toledo et al., 2013). Lack of ATR activity leading to unrestrained origin firing causes abnormally elevated formation of RPA-ssDNA which, in turn, engenders progressive exhaustion of the available nuclear pool of RPA and eventual formation of lethal DSB at RF in a phenomenon termed “replication catastrophe” (Toledo *et al*, 2017). Moreover, as RPA is also strictly required for NER (He et al., 1995), conditions that promote inordinate sequestration of RPA at stalled RF and/or at aberrantly activated replication origins post-UV were shown by our lab and others to cause S phase-specific defects in the removal of UV-induced DNA photoproducts (Auclair et al., 2008; Bélanger et al., 2018, 2015; Tsaalbi-Shtylik et al., 2014).

Several mechanisms have been shown to generate ssDNA in response to replicative stress and DNA damage: (1) During S phase, blockage of DNA polymerases causes their uncoupling from the MCM replicative helicase which continues to unwind DNA ahead of the stalled RF, resulting in abnormally large tracts of ssDNA (Byun et al., 2005). (2) Formation of reversed RF (Zellweger et al., 2015) creates nascent DNA ends that can be substrates for degradation by nucleases, e.g., MRE11 and EXO1, thereby generating ssDNA (Mijic et al., 2017). Unchecked nascent DNA degradation, termed “replication fork protection defect”, is prevented by several replicative stress response factors, including Rad51 and BRCA1/2 (Kolinjivadi et al., 2017b). (3) Defects in RF reversal or lesion bypass, e.g., TLS, can increase usage of PRIMPOL-dependent repriming downstream of the lesion, which generates ssDNA “gaps” behind replication forks (Quinet et al., 2021). (4) Following UV exposure, excision of lesion-containing oligonucleotides during NER transiently generates short stretches of ssDNA, which can be extended by the EXO1 nuclease to promote ATR activation (Giannattasio et al., 2010).

Given the demonstrated importance of adequate RPA availability in preventing the collapse of stalled RF (Toledo et al., 2017, 2013), mechanisms that limit ssDNA accumulation during exposure to replication-blocking genotoxins are expected to be major determinants of genomic stability. Here, we identify genetic networks governing RPA recruitment to DNA after UV irradiation using genome-wide CRISPR-Cas9 screening. Our data highlight a heretofore unknown role for SCAI, a factor previously implicated in gene transcription and DSB repair, in modulating the cellular response to UV-induced replication stress.

## RESULTS

### A genome-wide screen identifies regulators of RPA accumulation on DNA in response to UV irradiation

We sought to identify gene networks that restrict RPA accumulation on DNA during genotoxin-induced replicative stress. To this end, we optimized an existing method coupling flow cytometry, stringent washes, and immunofluorescence to measure ssDNA-associated (as opposed to free) RPA32 (one of the three subunits of the RPA complex) in U-2 OS human osteosarcoma cells in response to 254 nm UV (hereafter UV; Figure 1A) (Forment and Jackson, 2015). Exposure to 1 J/m^2^ UV caused detectable RPA recruitment to DNA at 1 and 3 h post-UV, which was largely resolved by 6 h (Figure 1A-B). In contrast, higher UV doses (3 or 5 J/m^2^) led to persistent accumulation of RPA (close to signal saturation) at all time points post-UV that we tested (Figure 1A-B). The dynamic range of this assay, within a 6-hour window, is therefore much larger at low (1 J/m^2^) vs higher doses of UV in U-2 OS cells (Figure 1B). As proof of principle for our experimental conditions, we treated cells with VE-821, a pharmacological ATR inhibitor which derepresses replication origins post-UV thereby generating abundant ssDNA (Toledo et al., 2013). As expected, ATR inhibition caused a strong increase in DNA-associated RPA in response to 1 J/m^2^ UV (Figure 1C-D).

**Figure 1:**
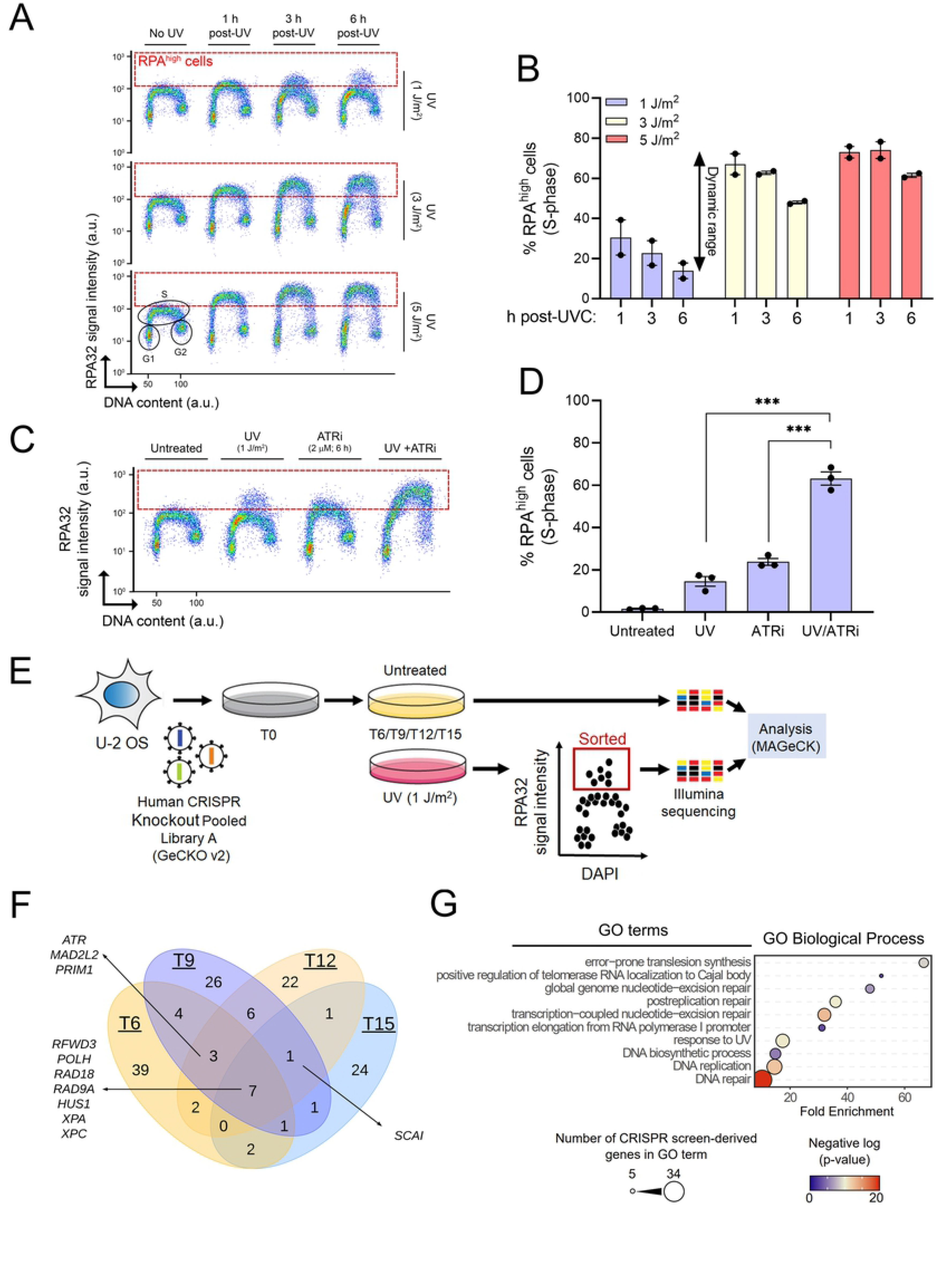
A flow cytometry-based CRISPR screen used to identify regulators of RPA-bound ssDNA formation. **A)** Immunofluorescence flow cytometry was used to measure ssDNA-RPA32 (y axis) and total DNA content (x axis; DAPI signal). Cells were treated with 1, 3 or 5 J/m^2^ UV or mock-treated and samples were collected 1, 3 or 6 h post-UV. The dashed red box identifies DNA-bound RPA^high^ cells. **B)** Quantification from (A). Values are the mean ± SEM from 2 independent experiments. **C)** Cells were mock treated or irradiated with 1 J/m^2^ UV +/- 2 µM of VE-821 (ATR inhibitor). Samples were harvested 6 h post-treatment. **D)** Quantification from (C). Values are mean ± SEM from 3 experiments. ***: p < 0.001. **E)** Schematic overview of the FACS-based CRISPR-Cas9 screen. Cells were irradiated with 1 J/m^2^ UV at 6-, 9-, 12- and 15-days post-infection with the GeCKOv2 lentiviral library (see material and methods). At each timepoint, mock-treated cells were collected to assess sgRNA representation. **F)** Venn diagram of the distribution of the genes recovered at each time point. **G)** Gene Ontology (GO) term enrichment analysis of genes identified in all the timepoints. Statistics in Figure 1: Student t-test.

We devised a CRISPR-Cas9 screening strategy employing the genome-wide GeCKOv2 lentiviral library (Sanjana et al., 2014; Shalem et al., 2014) in conjunction with the above-described flow cytometry assay (Figure 1E). U-2 OS cells were infected with the GeCKOv2 library and propagated for periods of 6, 9, 12, or 15 days to allow phenotypic expression. Cells were then either exposed to 1 J/m^2^ UV, or mock-treated. At 6 h post-UV, cells were fixed and labeled with anti-RPA32 antibodies followed by FACS to sort RPA^high^ cells (i.e., within the red dotted rectangle in Figure 1A, C). Following extraction of DNA from untreated and RPA^high^ cells, barcode sequences were amplified by PCR, and corresponding guide RNAs (sgRNA) identified by high-throughput sequencing. Results were then analyzed using the MAGeCK pipeline to identify sgRNA that are over-represented in the RPA^high^ population vs untreated controls (Li et al., 2014; Wang et al., 2019).

We found that sgRNA associated with the RPA^high^ fraction changed from day 6 to day 15 (Figure 1F, Supplementary Table S1), likely reflecting loss of sgRNA targeting essential and growth-promoting genes from the cell populations. Nevertheless, several genes were recovered at more than one time point (Figure 1F). Seven genes recovered at every time point encode factors with previously documented roles in the response to replicative stress and/or UV-induced DNA damage, as follows: RFWD3, a ubiquitin ligase that regulates both TLS and RPA recruitment to stalled replication forks (Elia et al., 2015; Gallina et al., 2021); DNA polymerase eta, a TLS polymerase that mediates accurate bypass of UV-induced CPD (Goodman and Woodgate, 2013), RAD18, a PCNA ubiquitin ligase involved in DNA damage tolerance (Branzei et al., 2008), RAD9, a component of the intra S phase checkpoint 911 complex (Parrilla-Castellar et al., 2004), and the NER pathway proteins XPA and XPC (Costa et al., 2003). Gene Ontology (GO-term) analysis of genes identified in our screen returned terms related to known pathways influencing the cellular response to UV-induced replicative stress, including error-prone translesion synthesis, nucleotide excision repair, DNA replication, and post-replication repair (Figure 1G).

We next evaluated siRNA-mediated depletion of individual “hits” from our screen on ssDNA-RPA formation post-UV. Genes from various functional groups were selected (Figure 2A). As expected, knockdown of RAD18, POLH, and XPC caused elevated ssDNA-RPA post-UV (Figure 2B-C). Our screen also identified factors whose potential roles in the UV-induced replicative stress response are incompletely characterized (Figure 2B-C): i) the TriC chaperonin complex (CCT2 and CCT8 subunits) which possesses several DNA repair/replication proteins as substrates (Yam et al., 2008), ii) the RUVBL1 chromatin remodeler, recently suggested to play roles in modulating the replicative stress response (Hristova et al., 2020), and iii) RIF1, a DNA double-strand break (DSB) repair factor that also regulates DNA replication origin activity (Hiraga et al., 2017; Zimmermann and de Lange, 2014). We note that downregulation of the above factors caused elevation in RPA-ssDNA specifically in S phase cells, consistent with the notion that most of the genes recovered in our screen act by mitigating replicative stress. We note that siRNA against RIF1 caused elevated RPA-ssDNA in the absence of UV, which might reflect the role of this gene in negatively regulating the activation of DNA replication origins in unperturbed cells (Hiraga et al., 2017). Overall, the results indicate that our screening strategy is competent in identifying mediators of the UV-induced DNA replication stress response.

**Figure 2:**
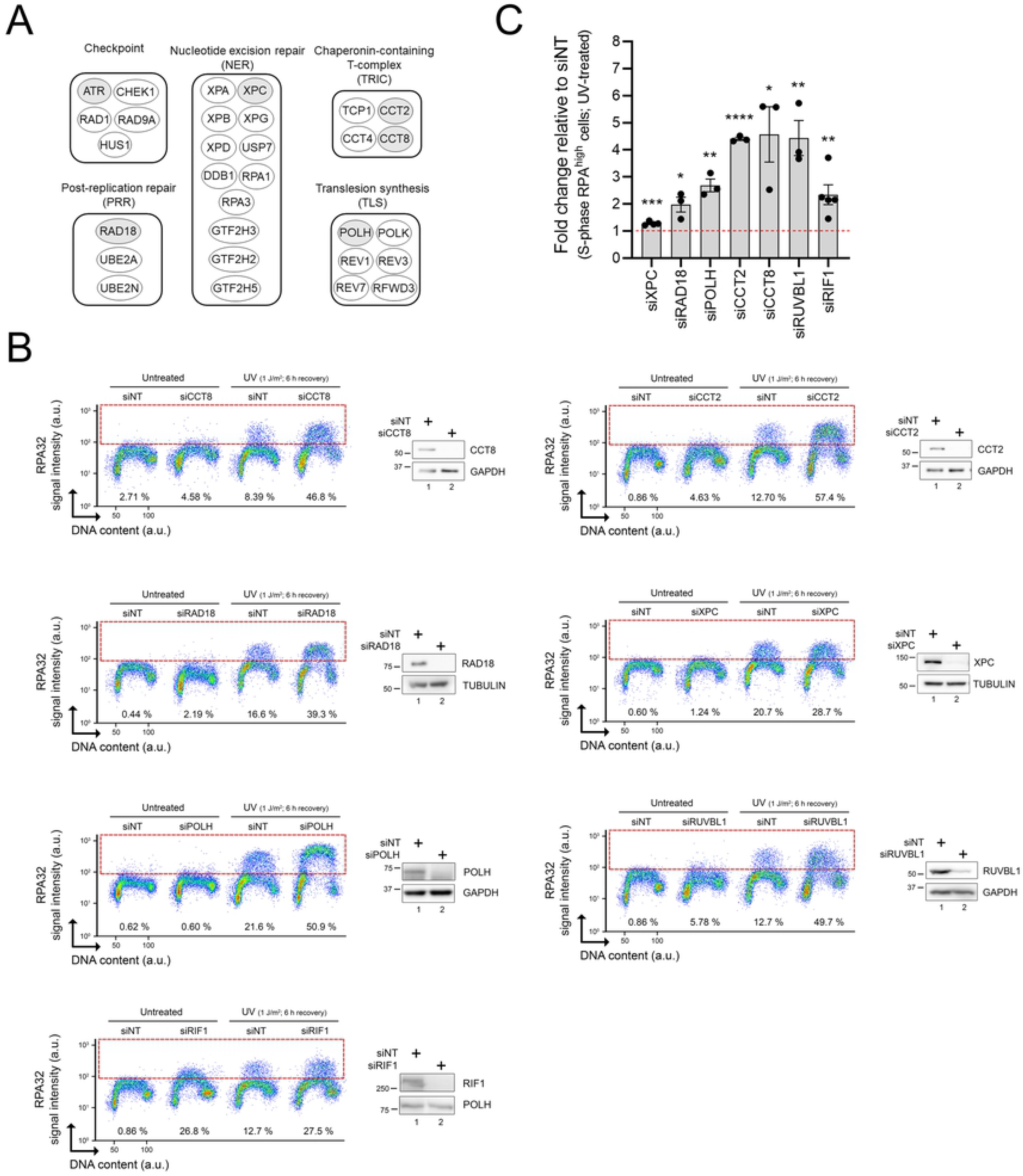
Validation of selected genes identified in the CRISPR-Cas9 screen. **A)** Main functional groups derived from genes recovered in the screen. Genes selected for further validation are shaded in grey. **B)** Representative immunofluorescence flow cytometry assays after siRNA-mediated depletion of selected genes. Cells transfected with non-targeting or gene-specific siRNAs were mock- or UV-treated (1 J/m^2^). % RPA^high^ cells (dashed box) were assessed 6 h after irradiation. Representative immunoblots and quantification of at least 3 independent flow cytometry experiments are shown. Values represent the mean ± SEM. *: p < 0.05, **: p < 0.01, ***: p < 0.001, ****: p < 0.0001. Statistics in Figure 2: Student t-test.

### SCAI is a novel regulator of the replicative stress response

The SCAI gene was recovered at multiple time points in our RPA-ssDNA screen (Figure 1F). SCAI has been reported to interact with 53BP1 to modulate DSB repair (Hansen et al., 2016; Isobe et al., 2017), and also to influence gene transcription (Brandt et al., 2009). However, any effect of SCAI on the response to genotoxin-induced replicative stress was unknown. We found that U-2 OS cells in which SCAI is either knocked-out via CRISPR-Cas9, or downregulated using siRNA, exhibited elevated RPA-ssDNA post-UV as compared to control cells (Figure 3A-D). Importantly, siRNA-mediated SCAI depletion also caused a similar phenotype in TOV-21G ovarian cancer cells (Supplementary Figure 1A-B). Like other genes identified in our screen, accumulation of RPA on DNA was observed primarily during S phase in cells lacking SCAI (Figure 3B-C), suggesting that this factor might modulate the response to replicative stress. Consistent with the elevated formation of RPA-ssDNA observed in Figure 3A-D, native immunofluorescence of incorporated BrdU, representative of ssDNA accumulation, was elevated in SCAI-depleted S phase cells post-UV as compared to control cells (Figure 3E). Exposure to other replicative stress-inducing drugs, e.g., cisplatin (CDDP) and 4-NQO, was also found to elevate RPA-ssDNA during S phase in SCAI-null compared to control cells (Figure 3F). Finally, our results indicate that SCAI-null U-2 OS cells are sensitized to UV and CDDP (Figure 3G-H). Taken together, these data show that upon exposure to genotoxins that cause replicative stress, SCAI acts to alleviate i) abnormal accumulation of RPA-ssDNA in S phase cells, and ii) loss of cell viability and/or reduced proliferation.

**Figure 3:**
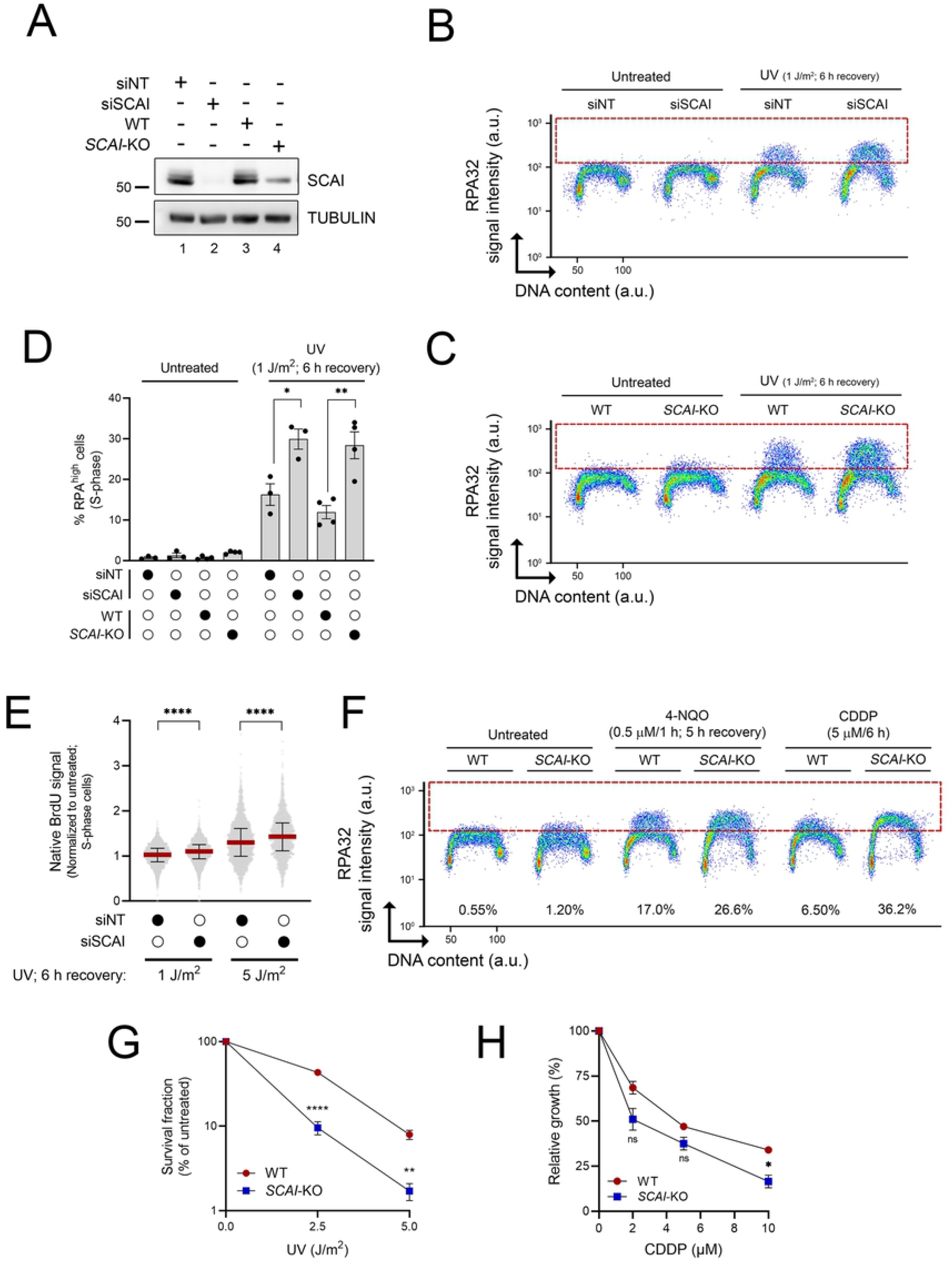
SCAI influences the replication stress response post-UV. **A)** Immunoblot of U-2 OS whole-cell extracts. **B-C)** Immunofluorescence flow cytometry measurements of DNA-associated RPA32 in control and SCAI-depleted cells 6 h after UV irradiation in U-2 OS. **D)** Quantification from (B-C). Values are the mean ± SEM from at least 3 independent experiments. *: p < 0.05, **: p < 0.01. **E**) Depletion of SCAI increases ssDNA generation post-UV. Control and SCAI-depleted cells were exposed to BrdU for 48 h, and then irradiated with UV as indicated. Native BrdU signal was assessed by immunofluorescence flow cytometry 6 h post-UV. Median is presented (red line), and error bars indicate the interquartile range. p-values were determined by Mann-Whitney test. ****: p < 0.0001 Mann-Whitney test. **F)** Immunofluorescence flow cytometry measurements of DNA-associated RPA32 in control and SCAI-depleted cells (as in B). WT and *SCAI*-KO cells were treated with 0.5 µM 4-NQO for 1 h and allowed to recover for 5 h or continuously exposed for 6 h to 5 µM cisplatin (CPPD). **G**) *SCAI*-KO cells are sensitive to UV as measured by clonogenic survival. Colonies were counted and normalized to untreated conditions. Histogram values are the mean ± SEM from 3 independent experiments. **: p < 0.01, ****: p < 0.0001. **H**) *SCAI*-KO cells are sensitive to CDDP. Cells were treated for 2h with CDDP in serum-free medium, followed by washing with PBS. Cells were then incubated in complete media for 3 days. Densitometry analysis of images of the stained dishes was used to evaluate cell growth. ns: non-significant, *: p < 0.05. Except for E, statistics in Figure 3 are performed using the Student t-test.

Several NER genes were recovered in our screen (Figures 1-2). Indeed, defective removal of UV-induced DNA lesions is expected to exacerbate RF stalling and accumulation of RPA-ssDNA in S phase cells. To address the possibility that SCAI regulates NER efficiency, we evaluated the DNA repair synthesis step of this pathway by quantifying incorporation of the nucleoside analog EdU in G1/G2 cells post-UV (Nakazawa et al., 2010; van den Heuvel et al., 2021). As expected, siRNA-mediated depletion of the essential NER factor XPC strongly attenuated repair synthesis compared to cells transfected with non-targeting siRNA (Figure 4A-B). In contrast, EdU incorporation post-UV was not reduced in SCAI-depleted vs control cells, suggesting that the global genomic NER subpathway is not compromised in the latter (Figure 4A-B). Similarly, we found that recovery of RNA synthesis post-UV as measured by incorporation of the nucleoside analog EU (Nakazawa et al., 2010; van den Heuvel et al., 2021), an indicator of the efficiency of the transcription-coupled NER subpathway, was similar in control vs SCAI-depleted cells but clearly defective in cells in which the essential NER factor XPA was knocked-down (Figure 4C-E). Overall, the above results indicate that lack of SCAI does not cause replicative stress by compromising NER-mediated removal of UV-induced DNA lesions.

**Figure 4:**
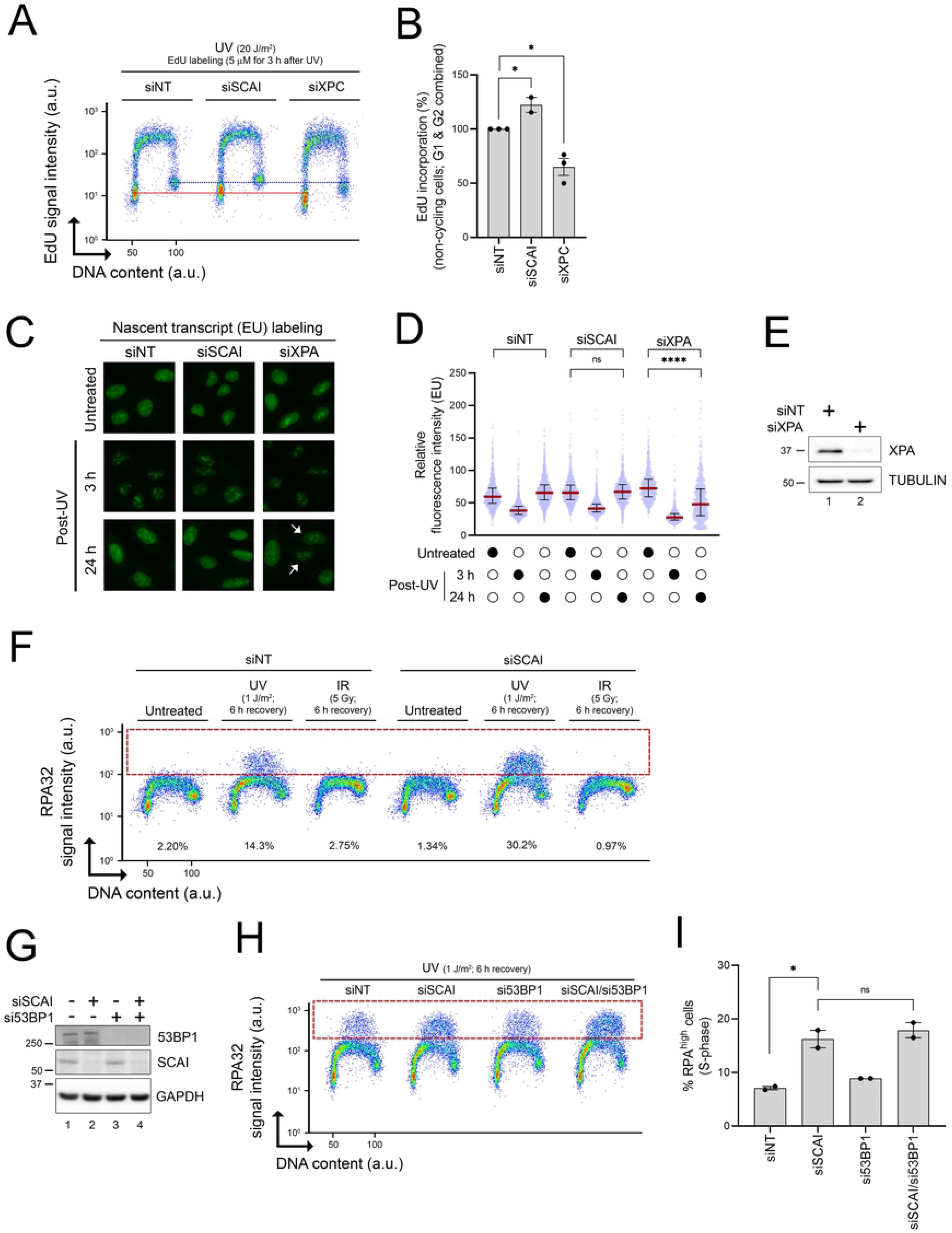
The functions of SCAI in the UV-induced replication stress response are unrelated to NER or 53BP1-dependent DSB repair. **A)** Flow cytometry was used to measure repair synthesis-associated EdU incorporation in G1/G2 cells (y axis). Total DNA content (x axis) was assessed with DAPI staining. Cells transfected with non-targeting (NT), SCAI-, or XPC-targeting siRNAs were irradiated with 20 J/m^2^ UV and allowed to recover for 3 h in medium containing 5 µM EdU. The red and blue dashed lines are positioned in the middle of the EdU signal of the G1 and G2 cell populations, respectively, of the siNT-treated cells to facilitate comparison. **B)** Quantification from (A). Histogram values are the mean ± SEM from at least 2 independent experiments and are relative to siNT-treated cells. *: p < 0.05. **C)** Representative images of 5-EU incorporation from cells transfected with the indicated siRNA. Cells were either mock- or UV-treated (6 J/m^2^) and samples collected 3 and 24 h after irradiation. White arrows indicate cells that did not incorporate 5-EU. **D)** Quantification from (C). Median is presented (red line), and error bars indicate the interquartile range. p-values were determined by Mann-Whitney test. ns: non-significant, ****: p < 0.0001. **E)** Validation of siRNA-mediated knockdown of XPA using immunoblot. **F)** Immunofluorescence flow cytometry was used to measure DNA-bound RPA32 (y axis) and DNA content (x axis; DAPI signal). Cells were irradiated with either 1 J/m^2^ UV or IR (5 Gy) and allowed to recover for 6 h prior to sample collection. The dashed red box identifies DNA-bound RPA^high^ cells. **G)** Immunoblot analysis from cells transfected with the indicated siRNA. **H)** siNT-, siSCAI-, si53BP1- and siSCAI/si53BP1-transfected cells were irradiated with 1 J/m^2^ UVC and allowed to recover for 6 h before immunofluorescence flow cytometry as in G. The dashed red box identifies DNA-bound RPA^high^ cells. **I)** Quantification from (I). Values represent the mean ± SEM from two independent experiments. ns: non-significant, *: p < 0.05. Statistics in Figure 4: Student t-test.

As mentioned previously, SCAI physically interacts with 53BP1 to modulate DSB repair (Hansen et al., 2016; Isobe et al., 2017). We therefore evaluated whether this functional interaction is relevant in the context of UV-induced RPA-ssDNA accumulation in S phase cells. Compared to the situation for UV, DSB-inducing ionizing radiation (IR) did not cause noticeable accumulation of RPA on DNA in either control or SCAI-depleted cells (Figure 4F), indicating that DSB processing, i.e., end resection, does not cause significant accumulation of RPA-ssDNA in S phase cells under our experimental conditions. We also found that depletion of 53BP1, alone or in combination with that of SCAI, did not influence levels of RPA-ssDNA post-UV in our assay (Figure 4G-I). Taken together, these data indicate that the abnormal response to replicative stress in cells lacking SCAI is unlikely to be related to defective 53BP1-dependent DSB repair.

### SCAI promotes DNA RF progression in UV-exposed cells

We next assessed the impact of SCAI on RF progression after UV irradiation using DNA fiber analysis. We found that both siRNA-mediated depletion and CRISPR-Cas9 knock-out of SCAI significantly compromised RF progression post-UV in U-2 OS cells (Figure 5A) as well as in two additional cancer cell lines: TOV-21G (ovarian cancer), and WM3248 (melanoma) (Supplementary Figure 1C-D). In contrast, SCAI depletion does not compromise RF progression in the absence of genotoxic treatment (Figure 5B). Importantly, our data also indicate that the negative impact of SCAI depletion on DNA RF progression after UV treatment is independent of 53BP1 (Figure 5C), as was the case for SCAI-dependent modulation of RPA-ssDNA levels (Figure 4H-J). Overall, these data demonstrate that SCAI influences RF progression after genotoxic stress.

**Figure 5.**
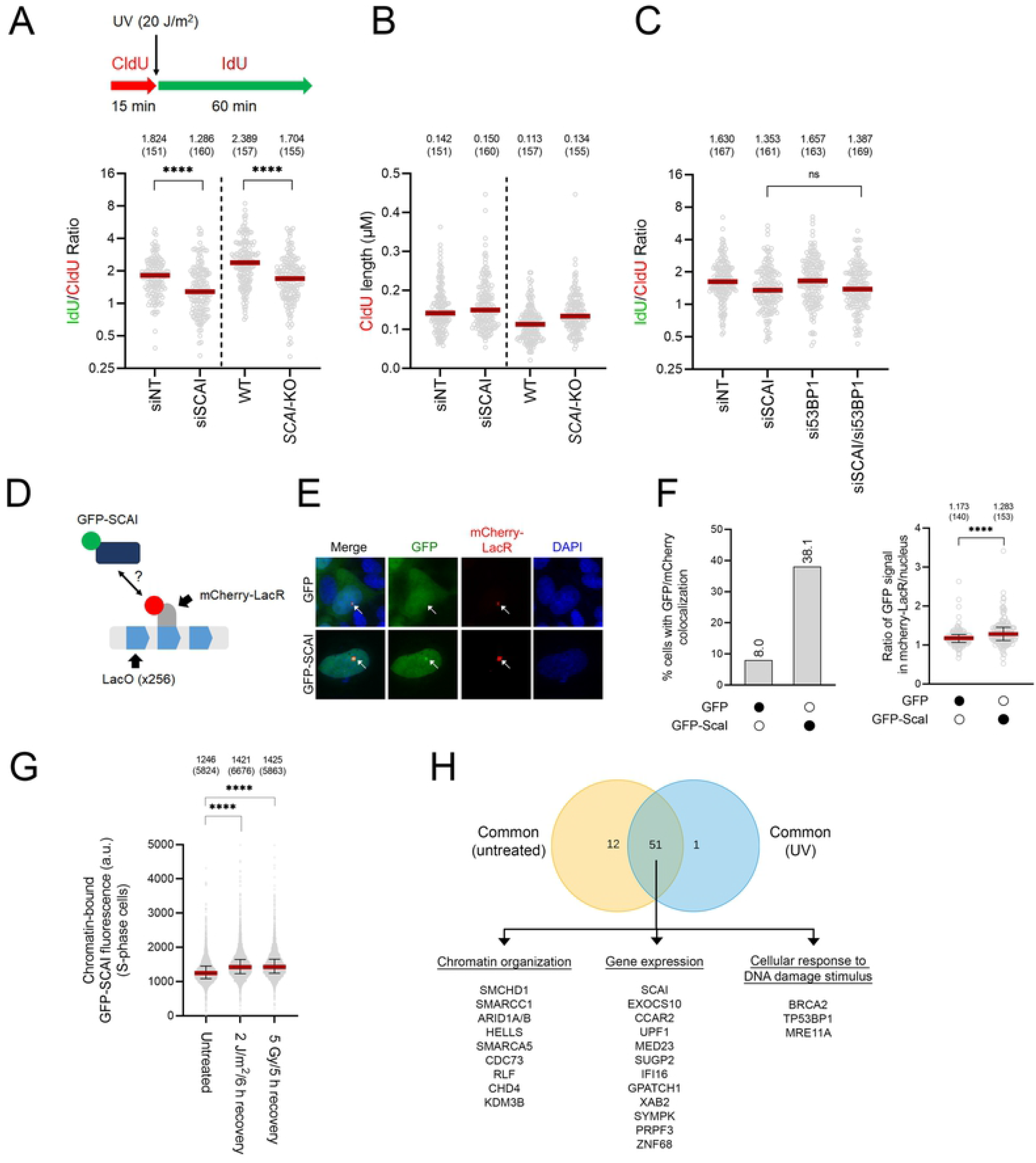
SCAI influences RF progression in cells exposed to UV. **A)** Cells lacking SCAI present defective RF progression post-UV. Cells were incubated with CldU (red) for 15 minutes, irradiated with UV (20 J/m^2^), and further incubated with IdU (green) for 60 minutes. A representative experiment is shown. Red bar: median. ****: p < 0.0001. **B)** Lack of SCAI compromises RF progression in the absence of genotoxic treatment. CldU fiber length from A are presented. Representative results are shown. Red bar: median. **C)** 53BP1 does not influence RF progression post-UV in cells lacking SCAI. Representative results are shown. Red bar: median. ns: non-significant, **: p < 0.01. **D)** Schematic of the assay used to evaluate recruitment of SCAI to stalled RF caused by binding of mCherry-LacR to a LacO array. **E)** Representative microscopy images for the assay described in D. **F)** SCAI is recruited to a mCherry-LacR-bound LacO array in the absence of DSB induction. Quantification of the experiment is presented in (D-E). Colocalization was scored positive when the GFP signal intensity in the mCherry-LacR foci was ≥ 1.5× the average nuclear GFP signal intensity. Results from a representative experiment are shown. ****: p < 0.0001. **G)** GFP-SCAI associates with DNA post-UV. Signal intensity was determined by flow cytometry +/- irradiation with 2 J/m^2^ UV or 5 Gy IR. Median is presented (red line), and error bars indicate the interquartile range. Cells were allowed to recover for 6 h (UV) or 5h (IR). a. u.: arbitrary units. ****: p < 0.0001. **H)** Proteins found in proximity of TurboID-SCAI after biotin labeling +/- UV (2 J/m^2^). Proteins from the untreated condition originate from 3 experiments, while proteins identified from UV-treated cells were identified from pooled results using cells allowed to recover for 1, 3 or 6 h after irradiation. The main Gene Ontology terms associated with proteins that overlap between UV-vs mock-treated are indicated. Statistics in Figure 5: Mann-Whitney test.

Biochemical purification of newly-replicated DNA using iPOND failed to identify SCAI as a component of stalled RF (Dungrawala et al., 2015). Nevertheless, it remained possible that interactions of SCAI with RF occur infrequently or are transient, thereby precluding detection of SCAI using this method. We therefore exploited a cell biology approach relying on the introduction of a 256XLacO array in U-2 OS cells expressing an mCherry-tagged LacR construct (Shanbhag et al., 2010). Recruitment of the LacR protein to the 256XLacOarray has previously been shown to be associated with RF stalling at this genomic region (Kim et al., 2020; Shanbhag et al., 2010). Cell lines harboring the 256XLacO array were engineered to express mCherry-LacR and either GFP-SCAI or control GFP. We confirmed that our GFP-SCAI fusion was functional by testing its previously reported ability to form nuclear foci in response to IR-induced DSB (Hansen et al., 2016; Isobe et al., 2017) (Supplementary Figure 2). Interestingly, GFP-SCAI colocalized frequently with the 256XLacO array compared to GFP, suggesting that SCAI is recruited in the vicinity of stalled RF *in vivo* (Figure 5D-F). Using fluorescence microscopy in cells subjected to stringent washes that remove proteins that are not bound to DNA, we found that UV elevates the binding of SCAI to DNA to a similar extent as IR (Figure 5G). Overall, the data suggest that replicative stress promotes recruitment of SCAI to DNA.

To further assess whether SCAI might be recruited in the vicinity of stalled RF, we used a variation of the BioID assay (TurboID) coupled to mass spectrometry, which permits rapid biotinylation, purification, and mass spectrometry-based identification of proteins in close spatial proximity to a protein of interest (Supplementary Table S2) (Cho et al., 2020; Roux et al., 2012). Consistent with a previous report indicating a role for SCAI in modulating transcription (Brandt et al., 2009), our analysis revealed that proteins involved in gene expression and chromatin organisation are biotinylated by TurboID-SCAI both in untreated and UV-exposed cells (Figure 5H). As expected, several peptides of the known SCAI-interacting DNA repair protein 53BP1 (Hansen et al., 2016; Isobe et al., 2017) were also recovered. Interestingly, BRCA2 and MRE11A, two homologous recombination proteins that are well-known to be recruited to, and to play important roles at, stalled RF (Kolinjivadi et al., 2017b) were identified as being in close physical proximity to SCAI in both UV- and mock-treated cells. Overall, our data support the notion that SCAI is recruited in the vicinity of RF in U-2 OS cells.

### EXO1 elevates RPA-ssDNA in the absence of SCAI

Several nucleases, including EXO1 and MRE11, act to generate ssDNA at stalled RF (Kolinjivadi et al., 2017b, 2017a; Lemaçon et al., 2017). Moreover, several recent reports indicate that replicative stress leads to the formation of unreplicated ssDNA gaps behind forks which can be extended by EXO1 and MRE11 (Cantor, 2021; Piberger et al., 2020; Quinet et al., 2021). We therefore tested whether these nucleases might promote RPA-ssDNA formation in cells lacking SCAI. Strikingly, we found that accumulation of DNA-bound RPA post-UV was completely abrogated upon siRNA-mediated depletion of EXO1 in SCAI KO cells, whereas the effect of MRE11 was more modest (Figure 6A-C). We therefore focused further characterization on the relationship between SCAI and EXO1-dependent DNA degradation, and found that EXO1 knockdown rescues UV-induced RF progression defects caused by lack of SCAI (Figure 6D). Based on the above, we reasoned that depletion of SCAI might favor EXO1-dependent nucleolytic degradation of nascent DNA at stalled RF (Lemaçon et al., 2017), leading to reduction in RF progression and to the accumulation of RPA-ssDNA. We found that cells lacking SCAI display modest nascent DNA instability compared to cells in which the well-known RF protection factor BRCA2 has been depleted (Figure 6E, H) (Mijic et al., 2017). Interestingly, co-depletion of both factors caused an additive effect with regard to either RF progression or RF protection (Figure 6E-F), suggesting that SCAI and BRCA2 might act via distinct mechanisms to protect stalled RF from nucleolytic degradation.

**Figure 6:**
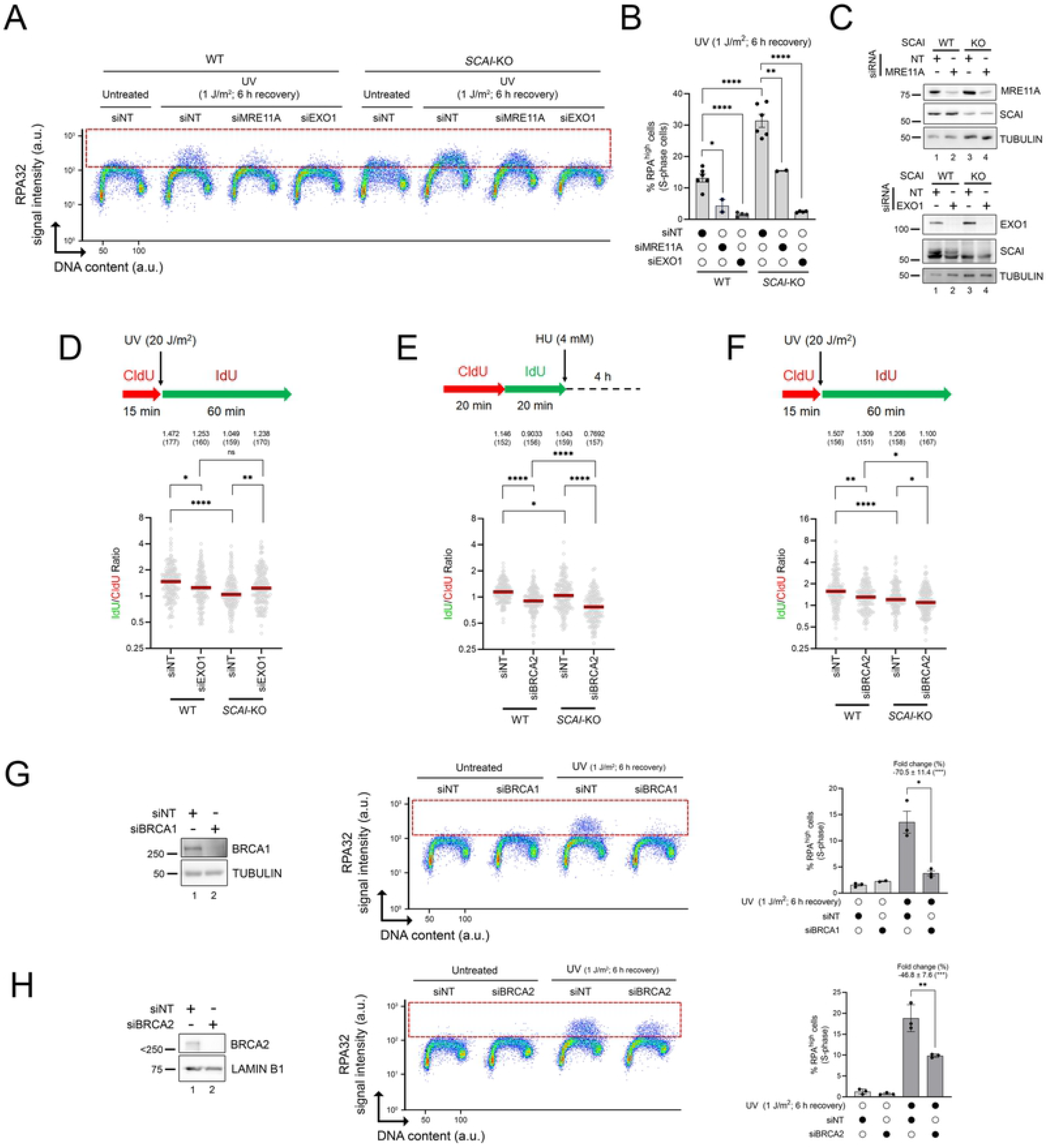
Accumulation of ssDNA-RPA depends on EXO1 in cells lacking SCAI. **A)** Depletion of EXO1, and to a lesser extent MRE11, rescues RPA-ssDNA accumulation in cells lacking SCAI post-UV. Cells were treated with 1 J/m^2^ UV or mock-treated and allowed to recover for 6 h. The dashed red box identifies DNA-bound RPA^high^ cells. **B)** Quantification from (A). Values represent the mean ± SEM from at least 2 independent experiments. *: p < 0.05, **: p < 0.01, ****: p < 0.0001. **C)** Immunoblot analysis showing EXO1 or MRE11 depletions in whole cell extracts from U-2 OS (WT) or *SCAI*-KO cells transfected with siRNAs. Tubulin serves as a loading control. **D)** Top: Schematic of the DNA fiber assay used to assess RF progression post-UV. Cells were incubated with CldU (red) for 15 minutes, irradiated with UVC (20 J/m^2^) and then incubated with IdU (green) for 60 minutes. Bottom: Dot plot of IdU/CldU ratio from WT (U-2 OS) and *SCAI* knockout cells transfected with siRNAs against EXO1. Red line: median. ns: non-significant, *: p < 0.05, **: p < 0.01, ****: p < 0.0001. A representative experiment is shown. **E)** Top: Schematic of the DNA fiber assay to monitor RF protection defects (nascent DNA degradation) after HU. Cells were incubated successively with CldU (red) and IdU (green) for 20 minutes each and then exposed to 4 mM HU for 4 h. Bottom: Dot plot of IdU/CldU ratio from WT (U-2 OS) and *SCAI* knockout cells transfected with siRNAs against BRCA2. Red line: median. *: p < 0.05, ****: p < 0.0001. A representative experiment is shown. **F)** Similar experiment as in (D) but from WT (U-2 OS) and *SCAI* knockout cells transfected with siRNAs against BRCA2. *: p < 0.05, **: p < 0.01, ****: p < 0.0001. **G-H)** Lack of BRCA1/2 does not cause RPA-ssDNA accumulation under our experimental conditions. Experiments were performed as in (A) but from U-2 OS (WT) cells transfected with siRNAs against BRCA1 (G) or BRCA2 (H). Values in rightmost panels are the mean ± SEM from at least 3 independent experiments. *: p < 0.05, **: p < 0.01. Tubulin and Lamin B1 serve as loading controls. Statistics in Figure 6: B, G, H: Student t-test. D-F: Mann-Whitney test.

We next tested directly whether RF protection defects, i.e., degradation of nascent DNA at reversed forks, contributes to the accumulation of RPA-ssDNA post-UV under our experimental conditions. BRCA1/2 are well-known to contribute to the protection of nascent DNA at stalled RF (Mijic et al., 2017; Schlacher et al., 2011). However, our screen did not identify BRCA1/2 (Supplementary Table S1), and moreover cells lacking either of these proteins did not display significant elevation of RPA-ssDNA in our assay (Figure 6G-H). In fact, depletion of either BRCA1 or BRCA2 (Figure 6G-H) led to a reduction in UV-induced ssDNA-RPA accumulation (Figure 6I-J). Taken together, the results suggest that nascent DNA degradation does not detectably contribute to UV-induced accumulation of RPA-ssDNA under our experimental conditions

EXO1 has been shown to extend ssDNA gaps left behind RF as a result of repriming and consequent replicative bypass of damaged DNA bases (Piberger et al., 2020). Such gap formation contributes to ssDNA generation in response to helix-destabilizing DNA adducts (Piberger et al., 2020). Previously published data also suggested that SCAI possesses the capacity to bind ssDNA (Hansen et al., 2016), raising the possibility that SCAI might directly influence the activity of EXO1 at ssDNA gaps. We purified SCAI and assessed its ability to bind various ssDNA-containing substrates *in vitro* (Figure 7A). Our data indicate that while SCAI readily binds ssDNA, this protein displays much lower affinity for dsDNA or a “splayed arms” DNA structure (Figure 7B, Supplementary Figure S3). Importantly, we found that incubation with SCAI significantly reduced EXO1 nucleolytic activity on a substrate containing a 34 base-long ssDNA gap *in vitro* (Figure 7C). Using S1 nuclease DNA fiber assays, we further found that the fold-change in size reduction of DNA due to S1 nuclease cleavage, which targets ssDNA gaps (Quinet et al., 2017), was unchanged in SCAI-depleted vs control cells (Figure 7D). This suggests that the frequency of ssDNA gap generation is similar in cells lacking SCAI compared to control. Finally, siRNA-mediated depletion of Primpol, an enzyme which promotes repriming and post-replicative gap formation after genotoxic stress (Quinet et al., 2021), strongly rescued ssDNA-RPA accumulation post-UV in SCAI-depleted cells (Figure 7E-F). Taken together, the above data suggest that in response to genotoxins that block RF progression, SCAI acts to limit EXO1-dependent nucleolytic extension of ssDNA gaps that are formed as a consequence of repriming and consequent lesion bypass.

**Figure 7:**
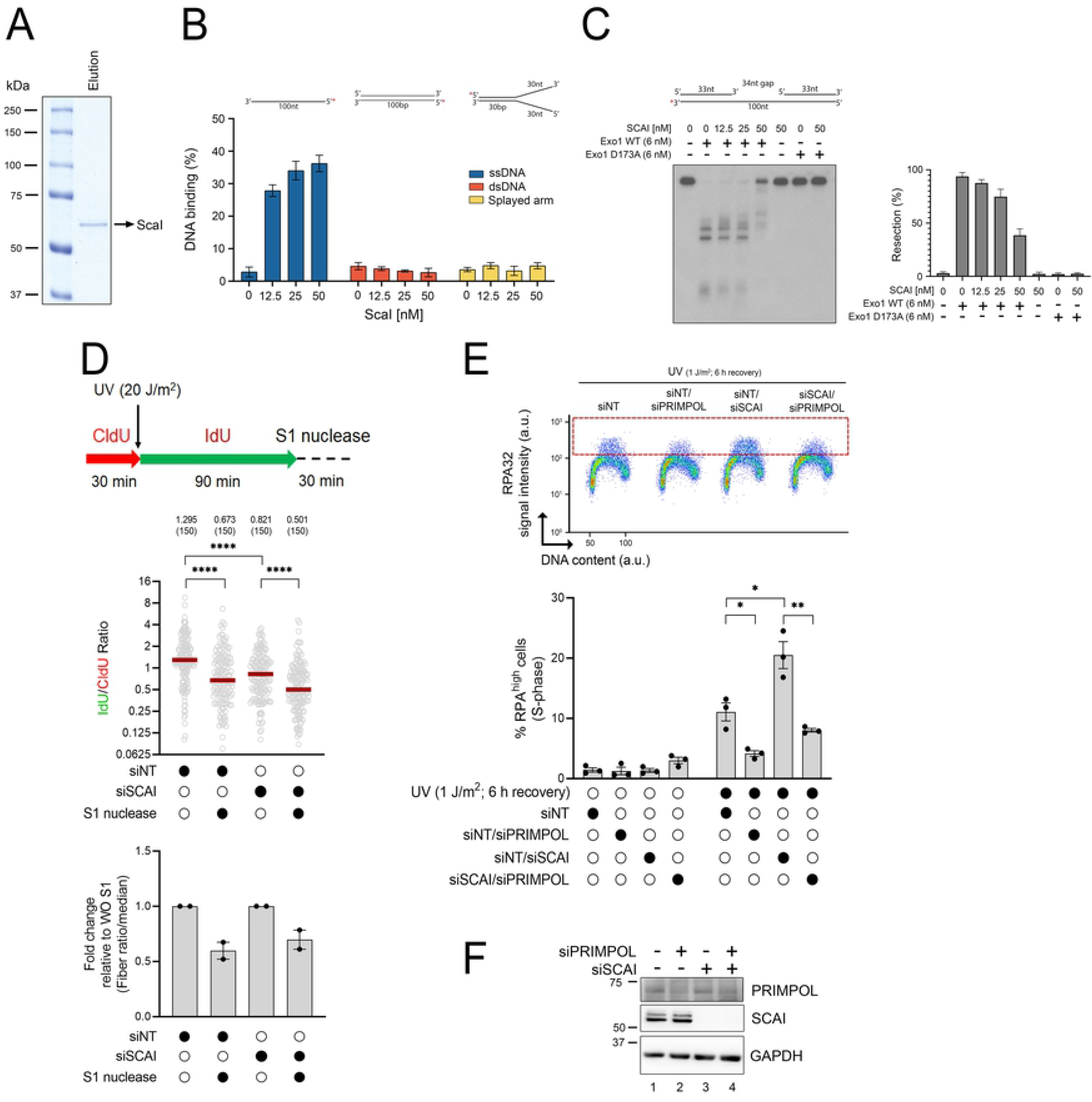
SCAI inhibits EXO1-mediated DNA resection. **A)** Recombinant SCAI protein was purified from insect cells, separated by SDS-PAGE and visualized by Coomassie blue staining. **B)** SCAI preferentially binds ssDNA over dsDNA. 5’- [^32^P]-labeled ssDNA, dsDNA, or “splayed arm” DNA were incubated with purified recombinant SCAI at increasing concentrations and the reaction products separated by acrylamide gel electrophoresis and visualized by autoradiography (See Supplementary Figure S3). Quantification of the percentage of SCAI-mediated DNA binding on ssDNA, dsDNA, and splayed arm substrates from 3 independent experiments. **C)** *In vitro* DNA resection assays using a 3’-[^32^P]-labeled “gapped” DNA substrate in the absence of any proteins, or with WT or a catalytically-inactive version of EXO1 (D173A) supplemented with purified recombinant SCAI. Quantification of the percentage of DNA resection from 3 independent experiments is shown. **D)** Depletion of SCAI does not increase ssDNA gap generation post-UV. Top: Schematic of the DNA fiber assay used to assess RF progression post-UV. Cells were incubated with CldU (red) for 30 minutes, irradiated with UV (20 J/m^2^) and then incubated with IdU (green) for 90 minutes. Cells were then treated or not with S1 nuclease. Bottom: Dot plot of IdU/CldU ratio from cells transfected with siRNAs as indicated. Red line: median. ns: non-significant, *: p < 0.05, **: p < 0.01, ****: p < 0.0001. **E)** Depletion of Primpol rescues ssDNA-RPA accumulation in cells lacking SCAI. Cells were transfected with the indicated siRNA, Cells were treated with 1 J/m^2^ UV or mock-treated. Cells were then allowed to recover for 6 h. The dashed red box identifies DNA-bound RPA^high^ cells. Histogram values represent the mean ± SEM from at least 2 independent experiments. *: p < 0.05, **: p < 0.01, ****: p < 0.0001. **F)** Validation of siRNA-mediated knockdown of PrimPol and SCAI by immunoblot.Statistics: Mann-Whitney test for D (DNA fiber dot plot), Student t-test for E.

## DISCUSSION

We developed a genome-wide screening strategy to identify genes limiting the formation of RPA-ssDNA in response to replication-blocking UV-induced DNA lesions. RPA-ssDNA serves as a platform for recruitment/activation of the intra-S phase checkpoint kinase ATR and other effectors of the replicative stress response (Iyer and Rhind, 2017; Maréchal and Zou, 2015). One important role of the ATR-mediated intra-S phase checkpoint is to limit the generation of RPA-ssDNA during genotoxin-induced replication stress by prohibiting origin activation. This, in turn, preserves adequate pools of RPA thereby forestalling genome-wide induction of DSB at persistently-stalled RF (Toledo et al., 2017, 2013). However, the precise molecular mechanisms underlying the formation of replication-associated DSB at stalled RF under conditions of limited RPA availability remain incompletely understood. ssDNA is known to be more susceptible to spontaneous cytosine deamination than dsDNA, leading to formation of abasic sites which may promote further replication fork stalling if left unrepaired (Lindahl, 1993). Moreover, ssDNA generated in the absence of ATR, which causes exhaustion of RPA pools, was found to be susceptible to cytosine deamination by APOBEC enzymes (Buisson et al., 2017). Finally, reducing the abundance of RPA stimulates the formation of secondary structures in ssDNA, which can lead to its nucleolytic degradation (Chen et al., 2013). The literature therefore clearly indicates that ssDNA is intrinsically less stable than dsDNA, and that its generation must be tightly controlled during replicative stress.

As expected, the ssDNA-RPA screen recovered several genes which, by virtue of their participation in the activation of the intra-S phase checkpoint, are important determinants of RPA-ssDNA generation. Indeed, this signalling cascade is known to limit the accumulation of RPA-ssDNA during replicative stress in several ways. As mentioned earlier, intra-S phase checkpoint signalling inhibits the initiation of new origins of replication, thereby restricting the number of stalled RF and consequent ssDNA formation (Santocanale and Diffley, 1998; Yekezare et al., 2013). Data from yeast also clearly demonstrate that intra S phase checkpoint mutants accumulate much longer stretches of ssDNA than wild type cells at individual stalled RF, although the precise mechanisms are not entirely clear (Sogo et al., 2002). Importantly, these stretches of ssDNA result at least in part from EXO1-dependent degradation of nascent DNA at stalled RF, which is inhibited by the intra S phase checkpoint kinase Rad53 in yeast (Segurado and Diffley, 2008).

As second category of “hits” from our RPA-ssDNA screen is involved in DNA damage tolerance via translesion synthesis (TLS). We previously demonstrated that lack of TLS polymerase eta, which is required for accurate bypass of UV-induced CPD, causes strong accumulation of RPA on DNA post-UV (Bélanger et al., 2015). Moreover, we and others showed that, as is the case for cells lacking intra S phase checkpoint signalling, ssDNA accumulation caused by defective TLS is sufficiently elevated to cause S phase-specific defects in UV-induced DNA photoproduct removal by sequestering RPA at stalled forks and preventing its action during NER (Auclair et al., 2010; Bélanger et al., 2015; Tsaalbi-Shtylik et al., 2014). Interestingly, recently published data indicate that defective TLS enhances the formation of post-replicative ssDNA gaps by favoring PRIMPOL-dependent repriming beyond damaged bases (Nayak et al., 2020; Quinet et al., 2021). Moreover, formation of such ssDNA gaps have been shown to cause strong sensitivity to replicative stress (Cong et al., 2021; Panzarino et al., 2021). It therefore seems likely that ssDNA gap formation behind RF underlie the strong representation of TLS polymerases, and regulators thereof, in our screen.

As expected, we also recovered genes encoding NER factors as regulators of RPA-ssDNA generation upon UV irradiation. NER-mediated removal of damaged DNA generates ssDNA gaps during the repair synthesis step in all phase of the cell cycle, which can be extended via the action of nucleases (Giannattasio et al., 2010). Nevertheless, the absence of NER activity presumably results in a larger number of persistent replication-blocking UV-induced lesions, leading to ssDNA formation specifically in S phase cells, which is what we observed (Figure 2B). We note however that the extent of RPA-ssDNA generation caused by NER defects was less pronounced than those caused by deficiencies in the intra S phase checkpoint or TLS pathways. This suggest that *i*) a large fraction of persistent UV-induced DNA lesions can be readily bypassed by DNA damage tolerance pathways during S phase, and consequently *ii*) that NER defects *per se* only cause modest elevation in replicative stress in human cells under our experimental conditions.

The RPA-ssDNA screen also identified several factors whose roles in modulating the cellular response to UV-induced replicative stress has not been as well documented compared with the above examples. TriC is a chaperone complex that assists in protein folding (Knowlton et al., 2021; Martín-Cófreces et al., 2021; Yam et al., 2008) and which has been reported to influence various cellular pathways including gene expression (Shaheen et al., 2021), cellular signalling (Weng et al., 2021), and protection against proteotoxic stress (Llamas et al., 2021). Interestingly, recent data indicate that activation of the integrated stress response, a cellular signalling cascade which responds to protein misfolding, leads to inhibition of histone gene synthesis and consequent formation of R-loops that are known to inhibit DNA RF progression (Choo et al., 2020). Curiously however, published data also show that inhibition of RF progression caused by lack of histone synthesis is not associated with dramatic elevation of RPA-ssDNA (Mejlvang et al., 2014). Since TriC assists in the folding of many proteins, the molecular mechanisms explaining its influence on DNA replication stress and RPA-ssDNA formation are likely complex, and their elucidation would require further experiments.

Rif1 plays several roles which might allow this factor to limit accumulation of RPA on DNA: *i*) regulation of DSB repair by interacting with the critical non-homologous end-joining factor 53BP1 (Chapman et al., 2013), *ii*) inhibiting origins of DNA replication by promoting dephosphorylation of the MCM complex (Hiraga et al., 2017; Mattarocci et al., 2014), and *iii*) preventing degradation of nascent DNA at stalled RF (Garzón et al., 2019). Our results indicate that defective 53BP1-dependent DSB repair does not cause an important accumulation of RPA-ssDNA in S phase cells. Furthermore, we showed that degradation of nascent DNA at reversed RF, i.e., defective RF protection, does not strongly contribute to RPA accumulation on DNA under our experimental conditions. We therefore speculate that, as is the case for cells lacking ATR (Toledo et al., 2013), abnormal activation of DNA replication origins probably contributes to elevated RPA-ssDNA generation in cells lacking Rif1. We note that our data are at odds with published reports indicating that cells lacking BRCA2, which are known to display strong fork protection defects, generate elevated ssDNA in response to HU (Duan et al., 2020). While the source of this discrepancy is unknown, it is possible that degradation of nascent DNA in response to UV only causes a modest amount of ssDNA which cannot be readily detected under our experimental conditions. We also note that our data is consistent with the fact that lack of BRCA1, which is well-known to cause severe RF protection defects, does not elicit S phase-specific NER defects due to sequestration of RPA at stalled RF (Bélanger et al., 2018).

We have identified SCAI as a new regulator of the replicative stress response in human cells. While our work was in preparation, another group used a different screening strategy to reveal that lack of SCAI sensitizes cells to cisplatin-induced DNA interstrand crosslinks (Adeyemi et al., 2021). This investigation also showed that SCAI protects nascent DNA at stalled RF against degradation by EXO1, thereby limiting the formation of ssDNA. Our work is generally consistent with these data, and moreover extends them by identifying SCAI as a regulator of DNA RF progression and ssDNA gap processing in response to UV-induced helix-destabilizing lesions. In addition, the REV3 subunit of TLS polymerase zeta was reported in the above-mentioned study to physically associate with SCAI to protect nascent DNA at stalled RF. This was found to be independent of REV7, the other pol zeta subunit, suggesting that pol zeta *per se* is not involved in restricting ssDNA accumulation during replicative stress. While we did not evaluate the role of the SCAI-REV3 interaction in response to UV, our screen did identify several TLS factors, including both REV3 and REV7, as negative regulators of ssDNA accumulation. Our data therefore suggest that in the context of UV-induced replicative stress, the role of REV3 as a subunit of pol zeta is probably important in preventing excessive generation of ssDNA. We also note that our *in vitro* data indicate that SCAI can act alone to limit EXO1 activity at ssDNA gaps, consistent with the notion that REV3 and SCAI may exert distinct roles during ssDNA gap processing.

Also consistent with the aforementioned recently published study (Adeyemi et al., 2021), we found that lack of SCAI leads to degradation of nascent DNA, i.e., RF protection defects, although the negative impact of SCAI depletion on nascent DNA stability is more modest than that caused by depletion of the well-known RF protection factor BRCA2. Nevertheless, depletion of EXO1 rescued UV-induced reduction in RF progression in cells lacking SCAI, suggesting that this latter protein promotes DNA replication by limiting degradation of nascent DNA at stalled RF. We also note that depletion of BRCA2 in SCAI-null cells caused additive defects in both RF progression and protection upon UV and HU, respectively, suggesting that these proteins act in a non-redundant manner to protect RF from nucleolytic activity. Since nascent DNA degradation at stalled RF in cells lacking BRCA2 does not cause significant accumulation of RPA-ssDNA post-UV under our experimental conditions, RF protection defects are unlikely to account for the observed ssDNA accumulation in SCAI KO cells. Interestingly, our *in vitro* experiment indicates that SCAI binds ssDNA with much greater affinity than either dsDNA, or splayed DNA junctions which resemble stalled RF. This is in agreement with published data showing interaction of SCAI with ssDNA, and with its co-localization with RPA in the context of DSB repair (Hansen et al., 2016). Moreover, we found that SCAI inhibits EXO1 activity on a ssDNA gap *in vitro*. Extension of ssDNA gaps by EXO1 and other nucleases has been shown to occur in response to lesions in template DNA (Piberger et al., 2020) and to significantly contribute to the formation of ssDNA upon replicative stress (Cong et al., 2021; Nayak et al., 2020; Panzarino et al., 2021). Taken together, the above leads us to propose that interaction between SCAI and ssDNA at post-replicative gaps might prevent nucleolytic extension of the latter by EXO1. While we did not formally investigate the impact of SCAI on EXO1-mediated degradation of nascent DNA at reversed forks, we note that this nuclease is known to act on both stalled RF and ssDNA gaps (Lemaçon et al., 2017; Piberger et al., 2020). It is therefore tempting to speculate that SCAI-dependent reduction of ssDNA formation at gaps or reversed RF might possess a similar mechanistic basis. Further experiments will be necessary to fully characterize the mechanisms through which SCAI impacts the generation of ssDNA in human cells.

## ACKNOWLEDGEMENTS

This work was supported by the following Canadian Institutes of Health Research (CIHR) grants: 201709PJT-388346 to H.W., 201603PJT-364096 to E.A.D., MOP-133442 to F.A.M. and FDN-388879 to J.Y.M. J.-F.L. is a recipient of a CIHR post-PhD fellowship. H.W. is the recipient of a Fonds de la recherche du Québec-Santé Senior scholarship. J.Y.M. is a Canada Research Chair in DNA repair and Cancer Therapeutics. Y.G. was supported by Fondation du CHU de Québec and FRQS PhD scholarships. D.A.R. and C.S. obtained FRQS PhD scholarships. C.S. received a PhD scholarship from the Cole Foundation. F.A.M. holds the Canada Research Chair in Epigenetics of Aging and Cancer. We thank Dr Juan Méndez (Spanish National Cancer Research Centre (CNIO)) for generously providing anti-PRIMPOL antibodies, and Dr Roger Greenberg (University of Pennsylvania) for the U-2 OS-265 cell line.

## MATERIAL AND METHODS

### Cell culture

U-2 OS and 293FT cells, purchased from ATCC and Invitrogen respectively, were cultured in Dulbecco’s Modified Eagle Medium (DMEM; Gibco/Thermo Fisher) supplemented with 10% fetal bovine serum (FBS; Wisent), 2 mM L-Glutamine (Gibco/Thermo Fisher), and antibiotics (100 U/mL penicillin and 100 μg/mL streptomycin; Gibco/Thermo Fisher). U-2 OS–FokI cells (also known as U-2 OS-265), obtained from Roger Greenberg (University of Pennsylvania) (Shanbhag et al., 2010), were cultured as above. U-2 OS Flp-In/T-Rex cells (hereafter U-2 OS FT), were cultured as above except for the addition of 100 µg/mL zeocin (InvivoGene) and 5 µg/mL blasticidin S (Gibco/Thermo Fisher) to the growth medium. Stable U-2 OS FT cell lines were maintained in the presence of 200 µg/mL hygromycin B (Gibco/Thermo Fisher) and 5 µg/mL blasticidin S. The ovarian cancer cell line TOV-21G (Provencher et al., 2000) was cultured in OSE medium (Wisent) supplemented with 10% FBS and antibiotics. The WM3248 human melanoma cell line (Coriel Institute) was propagated in Eagle’s MEM (Corning) containing 15% FBS, essential and nonessential amino acids (Corning), vitamins (Corning), L-glutamine, and antibiotics. All cell lines were cultured at 37°C under 5% CO_2_ in a humidified atmosphere. Cell lines were routinely tested for mycoplasma contamination by DAPI staining/fluorescence microscopy. All cell lines were authenticated by STR analysis (McGill University Genome Center).

### Generation of CRISPR-mediated knockout cell lines

U-2 OS CRISPR knockout lines were generated using the All-in-One plasmid encoding dual sgRNAs and fluorescent protein-coupled Cas9^D10A^ nickase (AIO-GFP; Addgene #74119) (Chiang et al., 2016). sgRNA pairs were designed using the WTSI Genome Editing online tool (http://www.sanger.ac.uk/htgt/wge/). AIO-GFP-containing sgRNA plasmids were transfected using Lipofectamine 2000 (Life Technologies/Thermo Fisher) as per manufacturer’s instructions. Two days later, transfected (EGFP-positive) cells were individually sorted by FACS into 96-well plates at a single-cell-per-well density for clonal expansion. Expanded clones were evaluated by immunoblotting to confirm knockdown of the protein of interest.

### Generation of stable inducible cell lines

U-2 OS Flp-In/T-REx cells were transfected using Lipofectamine LTX transfection reagent (Life Technologies/Thermo Fisher). Briefly, cells were seeded at 400 000 cells/well in a 6-well plate in 2 ml of complete DMEM without antibiotics. On day 1, cells were transfected with 100 ng of the pcDNA5-FRT/TO-based expression construct and 1 µg of pOG44 as per manufacturer’s instructions. On day 2 the transfected cells were transferred into a 10-cm dish in complete medium and, on day 3, selected by addition of blasticidin S and hygromycin B to the growth medium. The selection medium was changed every 3 days until visible colonies were observed. Colonies were then pooled, expanded, and protein expression monitored by Western blotting following the addition of 5 μg/mL doxycycline to the growth medium for a 24 h period.

### Genotoxic treatment

The following drugs were used in this study: ATRi: VE-821 (Selleckchem), cisplatin (CDDP) (Sigma), hydroxyurea (HU) (BioShop), 4-nitroquinoline 1-oxide (4-NQO) (Sigma). Treatment conditions are indicated in the corresponding figures. For 254-nm UV exposure, cell monolayers were washed with PBS, followed by irradiation in PBS with a Philips G25T8 germicidal lamp. The fluence was 0.2 J/m^2^/s, as monitored with a DCR-100X radiometer equipped with a DIX-254 sensor (Spectroline). Cells were exposed to IR using a ^137^Cs source (Gamma Cell 3000 Elan; Atomic Energy Canada) at a dose rate of 4.5×10^-2^ Gy/s.

### Clonogenic survival and growth assays

Cells were initially seeded at an appropriate density and, after attachment, washed with PBS and treated with various doses of UV in PBS. Following incubation for 14 days at 37°C, surviving colonies were stained with 0.5% methylene blue in 50% methanol. Colonies were counted and normalized to untreated samples to calculate relative survival. For CDDP sensitivity, 50 000 cells were seeded overnight in 6 cm dishes. CDDP was added for 2 h in serum-free medium, followed by washing with PBS. Cells were then incubated in complete media for 3 days. After staining with 0.5% methylene blue in 50% methanol, densitometry analysis was performed to assess cell growth (using Image J).

### Flow cytometry (FACS)

Protein bound to DNA were monitored by flow cytometry essentially as described (Forment and Jackson, 2015). Briefly, cells were harvested, washed once with PBS, and extracted in PBS-T buffer (0.2% Triton X-100 in PBS) to remove non-DNA-bound protein. Extracted cells were washed with PBS-B (PBS 1× + 1% BSA) and fixed in 2% formaldehyde for 30 minutes at room temperature. Cells were pelleted, washed with and resuspended in Perm/Wash buffer (BD Biosciences), and counted. Equal numbers of cells for each condition were incubated with primary antibody (1/100) in Perm/Wash buffer for 1 h at room temperature followed by incubation with Alexa Fluor-conjugated secondary antibody (1/200) in Perm/Wash for 30 minutes in the dark. Click-iT chemistry was then performed to identify S-phase cells, which had been labelled by adding 10 µM EdU to the cell culture medium 30 minutes before harvesting. Finally, cells were stained with DAPI and analyzed using an LSRII flow cytometer (BD Biosciences). The data were analyzed with FlowJo software (Flowjo LLC). Gates to assess enrichment of DNA-bound protein were established in untreated samples, and applied to all samples.

### siRNA transfection

For siRNA-mediated knockdown, cells were reverse-transfected with 20 nmol of siRNA using Lipofectamine RNAiMax (Thermo Fisher) as per manufacturer’s instructions. The medium was refreshed 24 h later and, unless otherwise stated, experiments were carried out at 72 h post-transfection. See Supplementary Table S3 for a list of siRNAs used in this study.

### Immunobloting

Whole cell extracts (WCE) were obtained by suspending cells in lysis buffer (25 mM Tris-HCl pH 7.5, 2% SDS). Lysed cells where heated for 5 minutes at 95 °C before being sonicated. Protein extracts were quantified with BCA reagent (Thermo Fisher) and analysed by SDS-PAGE. For immunoblotting, membranes were blocked in 5% milk/TBST (TBS + 0.1% Tween-20) and then incubated with primary antibody overnight at room temperature. Membranes were subsequently probed with secondary peroxidase-conjugated antibodies that had been incubated in 5% milk/TBST at room temperature for 1 h. ECL-based chemiluminescence was detected using an Azure c600 imager (Azure Biosystems). See Supplementary Table S3 for a list of antibodies used in this study.

### DNA fiber assay

DNA fiber assays were performed essentially as described (Quinet et al., 2017). Briefly, cells were sequentially labeled with two thymidine analogs, 30 μM 5-chloro-2’-deoxyuridine (CldU; Sigma-Aldrich) and 250 μM 5-iodo-2’-deoxyuridine (IdU; Sigma-Aldrich) for the indicated times. Labeled cells were loaded onto glass slides and lysed in spreading buffer (50 mM EDTA, 0.5% SDS and 200 mM Tris-HCl pH 7.4). DNA fiber tracks were obtained through DNA spreading and fixed in 3:1 methanol:acetic acid solution for 10 minutes. DNA fibers were then denatured in 2.5 M HCl for 80 minutes, blocked for 20 minutes in PBS containing 5% BSA at room temperature, and sequentially stained with primary antibodies against CldU (1:400, Abcam) and IdU (1:25, BD Biosciences) for 2 hr. This was followed by incubation with the corresponding secondary antibodies conjugated to various Alexa Fluor dyes for 1 h at room temperature. Lastly, slides were mounted with Immuno-Fluore (MP Biomedicals) and nascent DNA fibers visualized using a DeltaVision Elite microscope. At least 150 DNA fibers were counted per sample. Median values are shown (red line) in all figures.

### DNA fibers with S1 nuclease treatment

DNA fiber assays with ssDNA-specific S1 nuclease were performed as described immediately above with minor modifications. Cells were labeled with 30 µM CldU for 30 minutes, irradiated with UV and then labeled again with 250 µM IdU for 90 minutes. Cells were then permeabilized with CSK100 buffer (100 mM NaCl, 10 mM MOPS pH 7, 3 mM MgCl_2_, 300 mM sucrose and 0.5% Triton X-100) for 10 min at room temperature, treated with the S1 nuclease (Thermo Fisher) at 20 U/mL in S1 buffer (30 mM sodium acetate pH 4.6, 10 mM zinc acetate, 5% glycerol, 50 mM NaCl) for 30 minutes at 37°C, and collected by scraping in PBS-0.1% BSA. Nuclei were then pelleted at 7000 RPM for 5 minutes at 4°C. The supernatant was removed leaving the volume necessary to have a final concentration of 1500 nuclei/μl.

### Plasmids

Versions of SCAI tagged with either GFP or V5-TurboID were generated by LR cloning (Gateway) using pcDNA5-FRT-TO-eGFP (provided by Anne-Claude Gingras; University of Toronto) and pcDNA5-FRT-TO-V5-TurboID, respectively, as the destination vectors. All constructs were validated by DNA sequencing.

### CRISPR screen

The human <u>GE</u>nome-scale <u>C</u>RISPR <u>K</u>nock-<u>O</u>ut pooled library A (GeCKO v2) (Sanjana et al., 2014) was co-transfected into 293FT cells with the lentiviral packaging plasmids psPAX2 and pMD2.G (Addgene). Viral production was accomplished as described previously (Joung et al., 2017) with minor modifications. Briefly, 293FT cells were cultured in complete DMEM medium without antibiotics and seeded in T-225 flasks to achieve 80-90% confluence at the time of transfection one day later. 70 µl of PLUS reagent (Invitrogen) were diluted into 2.25 mL of Opti-MEM, briefly mixed, and incubated at room temperature for 5 minutes. Subsequently, DNAs from the following sources were added: 30.6 μg GeCKO pooled library A, 23.4 μg psPAX2, and 15.3 μg pMD2.G. Separately, 208 µl of Lipofectamine LTX was diluted in 4.5mL of Opti-MEM, and briefly mixed. The PLUS reagent/DNA and Lipofectamine LTX mixtures were then combined, gently inverted, and incubated at room temperature for 20 minutes. The combined mixture was carefully added to the T-225 flask. The medium was aspirated after 24 h and replaced with harvesting media (complete DMEM + 1% BSA). Viral supernatants were harvested 48- and 72-h post-transfection, combined, filtered through a 0.45µm Stericup filter unit (Milipore), concentrated 10X using the Lenti-X concentrator reagent, aliquoted, and frozen at −80 °C.

Transduction of U-2 OS cells with the sgRNA library was performed at an MOI of 0.3 to obtain 300× coverage. After puromycin selection, cells were maintained in exponential growth throughout the course of the CRISPR screen. To account for differences in protein depletion over time, monitoring of DNA-bound RPA^high^ cells by FACS after UV irradiation was carried out at 6-, 9-, 12- and 15-days post-transduction. For this purpose, 165 ×10^6^ transduced/puromycin-selected cells were seeded on fifteen 15-cm dishes 24 h prior to each timepoint. A control (unirradiated) dish was also included to facilitate discrimination of RPA^high^ cells. Cells were subsequently processed as described in the Flow Cytometery section above. Cells displaying enrichment of DNA-bound RPA (RPA^high^) were sorted using a FACSAria cell sorter (BD Biosciences).

Genomic DNA was extracted from sorted cells as described (Joung et al., 2017). DNA was also isolated from an aliquot of 19.5×10^6^ cells (=300× coverage) harvested at every experimental timepoint which serve as a means to address the sgRNA representation throughout the CRISPR screen time course. The genomic DNA concentration was measured by fluorimetry (Turner Biosystems) using the Quant-iT™ PicoGreen™ dsDNA Assay Kit (Thermo Fisher). sgRNA sequences were amplified by PCR with the NEBNext High Fidelity PCR Master mix using barcoded primers as described previously (Yau and Rana, 2018), before being subjected to next-generation sequencing on an Illumina NextSeq 550 appartus. Raw sequencing data were processed using Cutadapt to remove adaptors (Kechin et al., 2017) and trimmed to isolate 20-nt sgRNA sequences. The MAGeCK algorithm (Li et al., 2014) was used for sgRNA sequence quantitation, gene-level enrichment and ranking. Filtering criteria were further applied to the MAGeCK gene sets. Only genes with at least 2 positive sgRNA, displaying an RRA score lower than 1×10^-3^, with sgRNA read counts difference greater than 2 between representation (total) and sorted samples, and with ≥ 50 reads in the sorted sample, were considered for further analysis.

### Proximity labelling (TurboID)

Identification of the SCAI interaction network through spatial proteomics, using an N-terminally tagged SCAI (V5-TurboID-SCAI) construct generated by Gateway cloning from a sequence-validated entry vector, was performed essentially as described (Branon et al., 2018; Hesketh et al., 2017). Briefly, polyclonal populations of stable U-2 OS Flp-In/T-REx cells with integrated TurboID-SCAI were grown on 15-cm plates to 75% confluency (≈ 60 × 10^6^ cells). Bait expression was induced by addition to the growth medium of doxycycline (5 µg/mL) for 24 hr. Biotinylation *in vivo* of potential protein partners was accomplished for 1, 3 and 6 hr prior to the end of the 24 h bait expression period on UV-treated (2 J/m^2^) or mock-treated cells by the addition of 500 µM biotin to the medium. Cells were kept on ice, washed extensively with cold PBS, lysed, sonicated, and biotinylated proteins purified with streptavidin-sepharose beads. Proteins were directly converted into peptides using the on-beads digestion technique as described (Dubois et al., 2016). Mass spectrometry analysis was performed as described (Lambert et al., 2020). An arbitrary threshold was applied to remove common background contaminants from protein partners identified in the TurboID assay. Proteins with at least 20 peptides and present in less than 10% of BioID experiments as listed in the Crapome database (Mellacheruvu et al., 2013) were selected for further analysis. Biological processes associated with the trimmed hit list were analysed using PANTHER (Mi et al., 2019).

### SCAI recruitment to LacR foci

For monitoring recruitment of GFP-tagged SCAI to mCherry–LacR–NLS foci, 150 000 U-2 OS– FokI cells were seeded on glass coverslips in a 6-well plate without induction of FokI. Twenty-four hours later, cells were transfected using 1µg of pDEST-mCherry-LacR-NLS (provided by Xu-Dong Zhu; McMaster University) and 1 µg of pcDNA5-FRT-TO-(eGFP-SCAI). 48 h after transfection, cells were fixed with 4% methanol-free formaldehyde/2% sucrose for 15 minutes at room temperature, washed successively with PBS and CSK buffer (100 mM NaCl, 300 mM sucrose, 10 mM PIPES pH 6.8, 3 mM MgCl_2_), permeabilized with CSK-T buffer (CSK buffer + 0.5 % Triton X-100) and stained with DAPI.

### Unscheduled DNA synthesis assay

Unscheduled DNA synthesis (UDS) post-UV was monitored by flow cytometry. Briefly, cells were irradiated with UV (20 J/m^2^) and allowed to recover in complete media containing 1% FBS and 5 µM EdU for 3 h. EdU-labelled cells were then processed as described above in the Flow Cytometry section. To assess the relative intensity of EdU in non-cycling cells, a dumb channel was used to isolate G1 and G2 cell populations. The median value from each condition was determined and set to 100% for the non-targeting siRNA (siNT).

### RNA synthesis recovery (RSR) assay

Visualization of nascent transcription by 5-ethynyl-uridine (EU) labeling post UV was performed as described (van den Heuvel et al., 2021). Briefly, siRNA-transfected cells grown on coverslips were mock-treated or irradiated with UV (6 J/m^2^). Cells were allowed to recover for either 3 or 24 h, and pulse-labelled with 400 µM of EU for 1 h prior to harvesting. In all cases, 24 h prior to the RSR assay, cells were grown in complete media containing 1% FBS to favor incorporation of EU. Labeled cells were processed as described in the above Flow Cytometry section except that the concentration of Alexa647-azide was increased to 10 µM in the Click-iT reaction, and DAPI staining was performed in analysis buffer that does not contain RNase.

### Recruitment of SCAI to DSB

For monitoring recruitment of SCAI to IR-induced DSB, 400 000 U-2 OS Flp-In/T-REx cells expressing either GFP alone or GFP-SCAI were seeded on glass coverslips in a 6-well plate in doxyclycline-containing media to induce protein expression. 24 h after seeding, cells were mock- or IR-treated. Cells were fixed with 4% PFA/2% sucrose for 15 min at room temperature, washed with PBS, permeabilized with CSK buffer (100 mM NaCl, 300 mM sucrose, 10 mM PIPES pH 6.8, 3 mM MgCl_2_, 0.5% Triton-X-100), stained with DAPI, and mounted on microscopy slides for imaging.

### Native BrdU assay

Assessment of native BrdU levels by flow cytometry was performed as described (Tkáč et al., 2016). Briefly, siRNA-transfected U-2 OS cells were grown in 20 µM BrdU-containing media for 48 hr before being mock- or UV-treated.

### Recombinant SCAI Purification

SCAI was tagged at the N-terminus with GST and at the C-terminus with His10 and was expressed and purified in Sf9 insect cells by infection with baculovirus generated from a pFASTBAC plasmid according to the manufacturer’s instructions (Bac-to-Bac, ThermoFisher). Transfection of Sf9 cells was carried out using Cellfectin II reagent (Thermo Fisher). Sf9 cells were infected with the generated SCAI baculovirus. 72 h post-infection, cells were harvested by centrifugation and the pellet frozen on dry ice. Cells were lysed in Buffer 1 (1X PBS containing 150 mM NaCl, 1 mM EDTA, and 1 mM DTT) supplemented with 0.05% Triton X-100 and protease inhibitors. Cell lysates were incubated with 1 mM MgCl_2_ and 2.5 U/ml benzonase nuclease at 4°C for 1 h followed by centrifugation at 35000 rpm for 1 h. Soluble cell lysates were incubated with GST-Sepharose beads at 4°C with gentle rotation. Beads were washed twice with Buffer 1 followed by incubation with Buffer 2 (Buffer 1 with 5 mM ATP, 15 mM MgCl_2_). Sepharose GST beads were washed twice with Buffer 3 (1X PBS supplemented with 200 mM NaCl) and once with P5 Buffer (20 mM NaHPO_4_, 20 mM NaH_2_PO_4_, 500 mM NaCl, 10% glycerol, 0.05% Triton-X-100, 5 mM Imidazole) followed by cleavage with PreScission protease (60 U/ml, GE Healthcare Life Sciences). The beads were applied to a column and the elution was collected and completed to 10 mL with P5 Buffer. The eluate was then incubated with TALON beads (ClonTech). Beads were washed twice with P5 Buffer and once with P30 Buffer (P5 supplemented with 25 mM Imidazole). The beads were applied to a column and the proteins eluted twice using P500 Buffer (P5 supplemented with 495 mM Imidazole). Proteins were then dialyzed in Storage Buffer (20 mM Tris-HCl, pH 7.4, 200 mM NaCl, 10% glycerol, 1 mM DTT) and stored in aliquots at −80°C.

### SCAI DNA Binding Assay and DNA substrates

JYM696: GGGCGAATTGGGCCCGACGTCGCATGCTCCTCTAGACTCGAGGAATTCGGTACCCCGGGTTCGAAATCGATAAGCTTACAGTCTCCATTTAAAGGACAAG

JYM698: CTTGTCCTTTAAATGGAGACTGTAAGCTTATCGATTTCGAACCCGGGGTACCGAATTCCTCGAGTCTAGAGGAGCATGCGACGTCGGGCCCAATTCGCCC

JYM925: GGGTGAACCTGCAGGTGGGCAAAGATGTCCTAGCAATGTAATCGTCAAGCTTTATGC CGT

JYM926: ACGCTGCCGAATTCTACCAGTGCCAGCGACGGACATCTTTGCCCACCTGCAGGTTCA CCC

5’-end ^32^P-labelled DNA substrates were generated using T4 PNK (NEB) and [ɣ-^32^P]ATP (PerkinElmer). End labelled JYM696 was used as the ssDNA substrate. dsDNA was produced by annealing JYM698 with labelled JYM696, and splayed arm DNA generated by annealing labelled JYM925 with JYM926. Both substrates were purified by PAGE.

DNA binding assays: The DNA-binding reactions (10 μl) contained the indicated DNA substrates (100nM) and the indicated concentrations of purified SCAI in MOPS buffer (25 mM MOPS at pH 7.0, 60 mM KCl, 0.2% Tween-20, 2 mM DTT, and 5 mM MgCl2). Reaction mixtures were incubated at 37°C for 15 minutes and transferred on ice. The reactions were subjected to electrophoresis on an 8% polyacrylamide gel at 4°C. Gels were dried for 35 minutes at 85°C on Whatman paper and visualized by autoradiography. Densitometric analyses were performed using a FLA-5100 phosphorimager (Fujifilm) and quantified using the Image Reader FLA-5000 v1.0 software.

### *In vitro* resection assays with SCAI and EXO1

JYM5735: AGAGGAGCATGCGACGTCGGGCCCAATTCGCCC

JYM5736: CTTGTCCTTTAAATGGAGACTGTAAGCTTATCG

3’-end ^32^P-labelled gapped DNA was generated using TdT (NEB) and [α-^32^P]dATP (PerkinElmer). The gapped DNA substrate was produced by annealing JYM5735 and JYM5736 oligonucleotides with ^32^P-labelled JYM696 and purified by PAGE. In vitro reactions were conducted using the gapped DNA probe in standard buffer (20 mM HEPES pH 7.5, 0.1 mM DTT, 0.05% Triton X-100, 100 µg/mL BSA) with 2 mM ATP and 5 mM MgCl_2_. Reactions were initiated on ice by adding the indicated concentrations of purified SCAI and transferred immediately to 37 °C for 5 minutes to allow binding of SCAI on the gapped DNA substrates. Subsequently, 6 nM purified EXO1 WT or EXO1 D173A (Exonuclease-dead) were added and transferred immediately to 37 °C for 30 minutes. Reactions were stopped by proteinase K treatment for 30 minutes at 37 °C. Products were analyzed on an 8% denaturing polyacrylamide/urea gel. Gels were dried for 2 h at 85°C on Whatman paper and visualized by autoradiography. Densitometric analyses were performed using a FLA-5100 phosphorimager (Fujifilm) and quantified using the Image Reader FLA-5000 v1.0 software.

### Statistics and reproductibility

For the DNA fiber experiments, the Mann-Whitney statistical test was used. For other assays, Student’s *t*-test (two-tailed) was used. Statistical analyses were performed using GraphPad Prism 9 software. Statistical significance is indicated for each graph (ns = not significant, for *p* > 0.05; * for *p* < 0.05; ** for *p* < 0.01; *** for *p* < 0.001; **** for *p* < 0.0001). All assays throughout this study were repeated at least twice.

### Image acquisition and analysis

Microscopy was performed using a DeltaVision fluorescence microscope equipped with SoftWorx (GE Healthcare). Images were analysed using a custom Python 3.6 script. Nuclei were segmented with DAPI staining channel images using Otsu’s thresholding, followed by extraction of the average fluorescence intensity per cell in the other channels. For Figure 5E-F, mCherry signal was thresholded using Otsu’s method to identify and segment the LacO array. Average GFP fluorescence was then calculated within this region and compared to the average of the total nuclear GFP value. GFP signals that were ≥ 1.5 fold higher than the average nuclear value were deemed as representing colocalization. In the case of DNA fiber assays, fiber length was measured manually using Image J.

---

**Figure S1:**
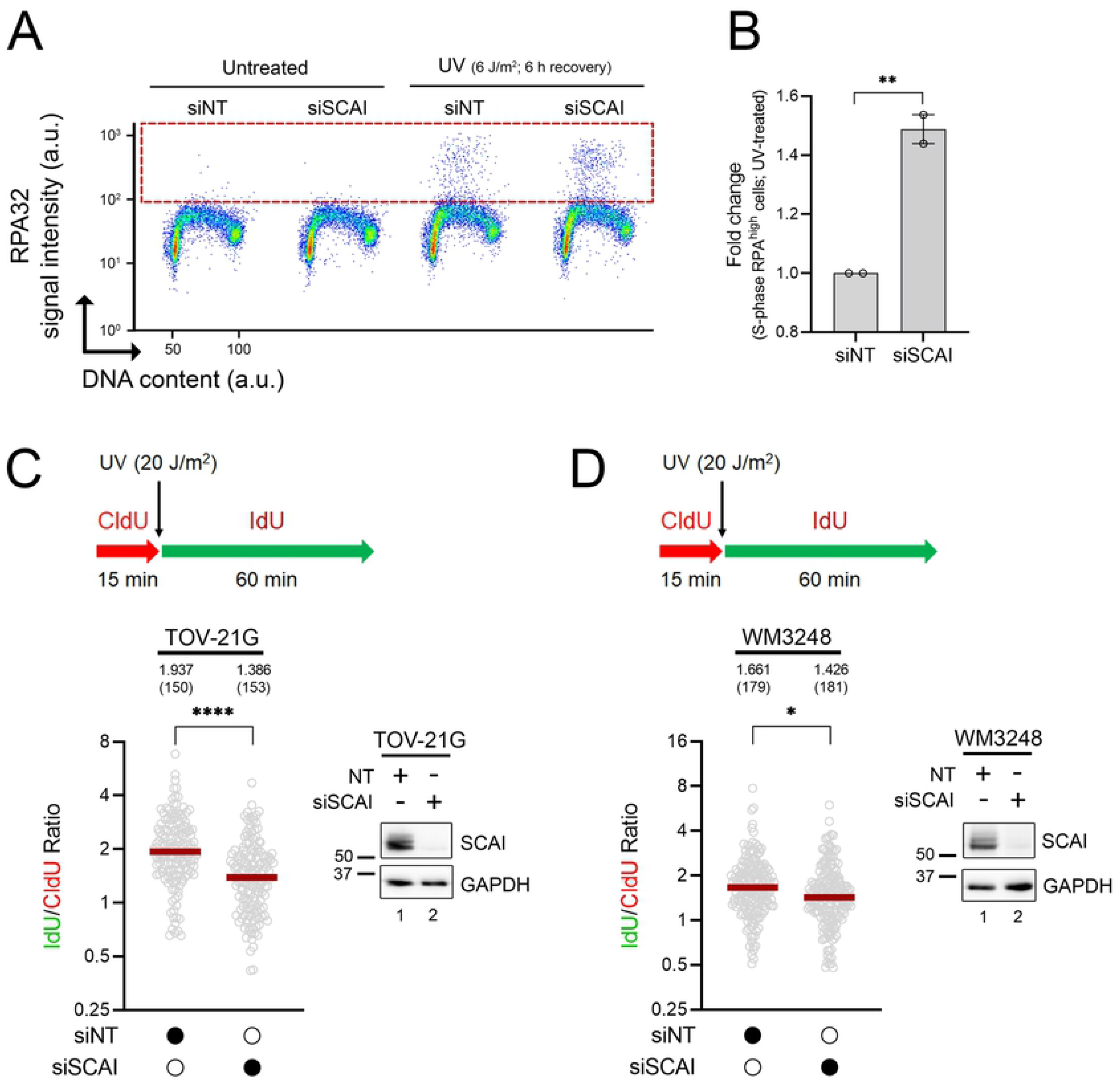
SCAI functions in the replication stress response in cellular backgrounds other than U-2 OS. **A)** Representative immunofluorescence flow cytometry assays after siRNA-mediated depletion of SCAI in TOV-21G cells. Cells were mock- or UV-treated (1 J/m^2^). % RPA^high^ cells (dashed box) were assessed 6 h after irradiation. **B)** Quantification from (A). **: p < 0.01, Student t-test. **C-D)** SCAI downregulation caused UV-induced reduction of RF progression in TOV-21G ovarian cancer (C) and WM3248 melanoma cell lines (D). Top: Schematic of the DNA fiber assay for fork progression assessment upon UV. Cells were first incubated with CldU (red) for 15 minutes, irradiated with UV (20 J/m^2^) and then incubated with IdU (green) for 60 minutes. Bottom: Dot plot of IdU/CldU ratio from siNT and siSCAI-transfected. Representative results from 2 independent experiments. Red line: median. *: p < 0.05, ****: p < 0.0001, Mann-Whitney test.

**Figure S2:**
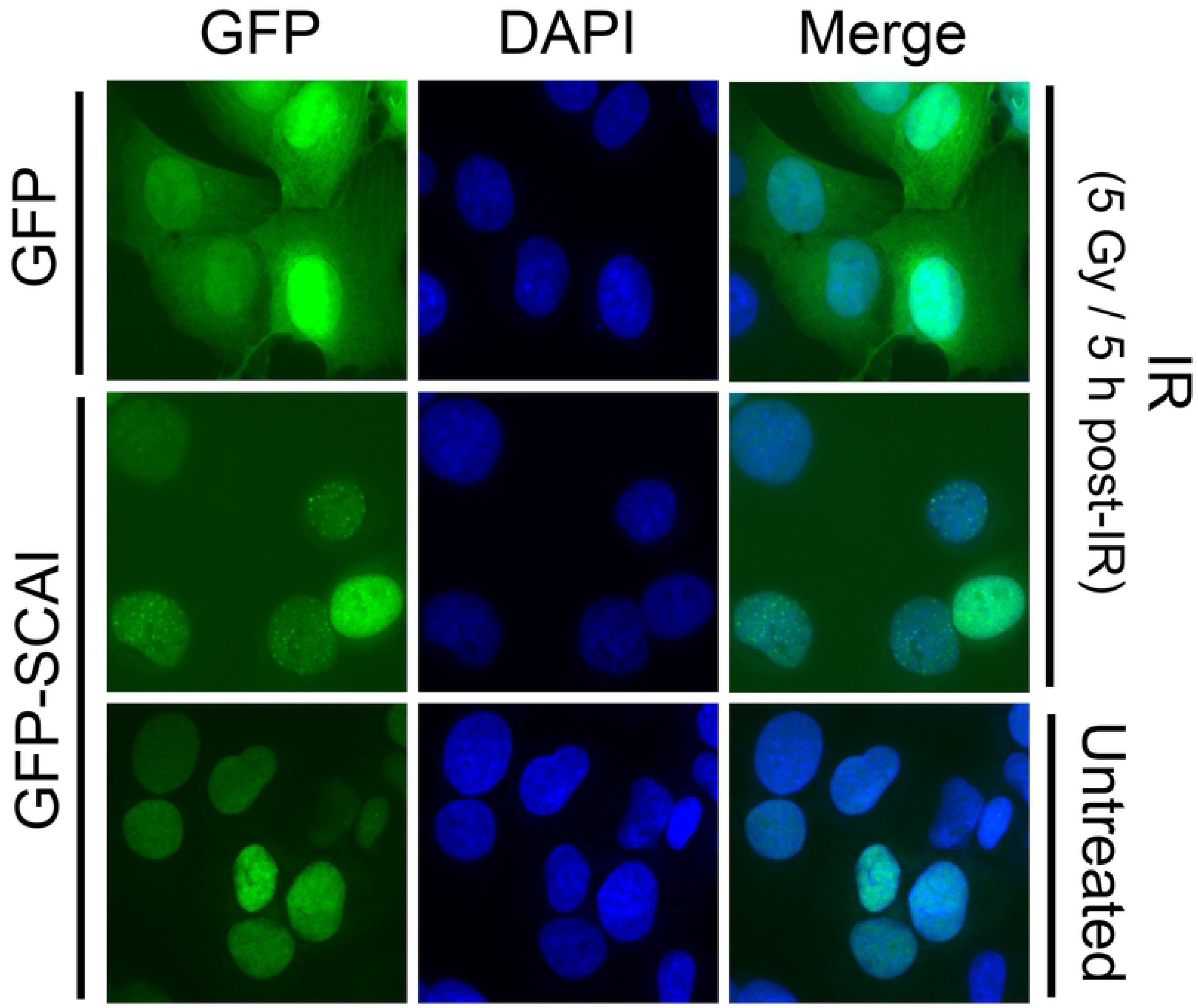
Functional validation of the GFP-SCAI construct. **A)** Recruitment of SCAI to IR-generated DSB repair foci. U-2 OS Flp-In/T-REx cells with a stably integrated GFP-SCAI construct were exposed to IR (5 Gy) and fixed/imaged after an incubation period of 5 h.

**Figure S3:**
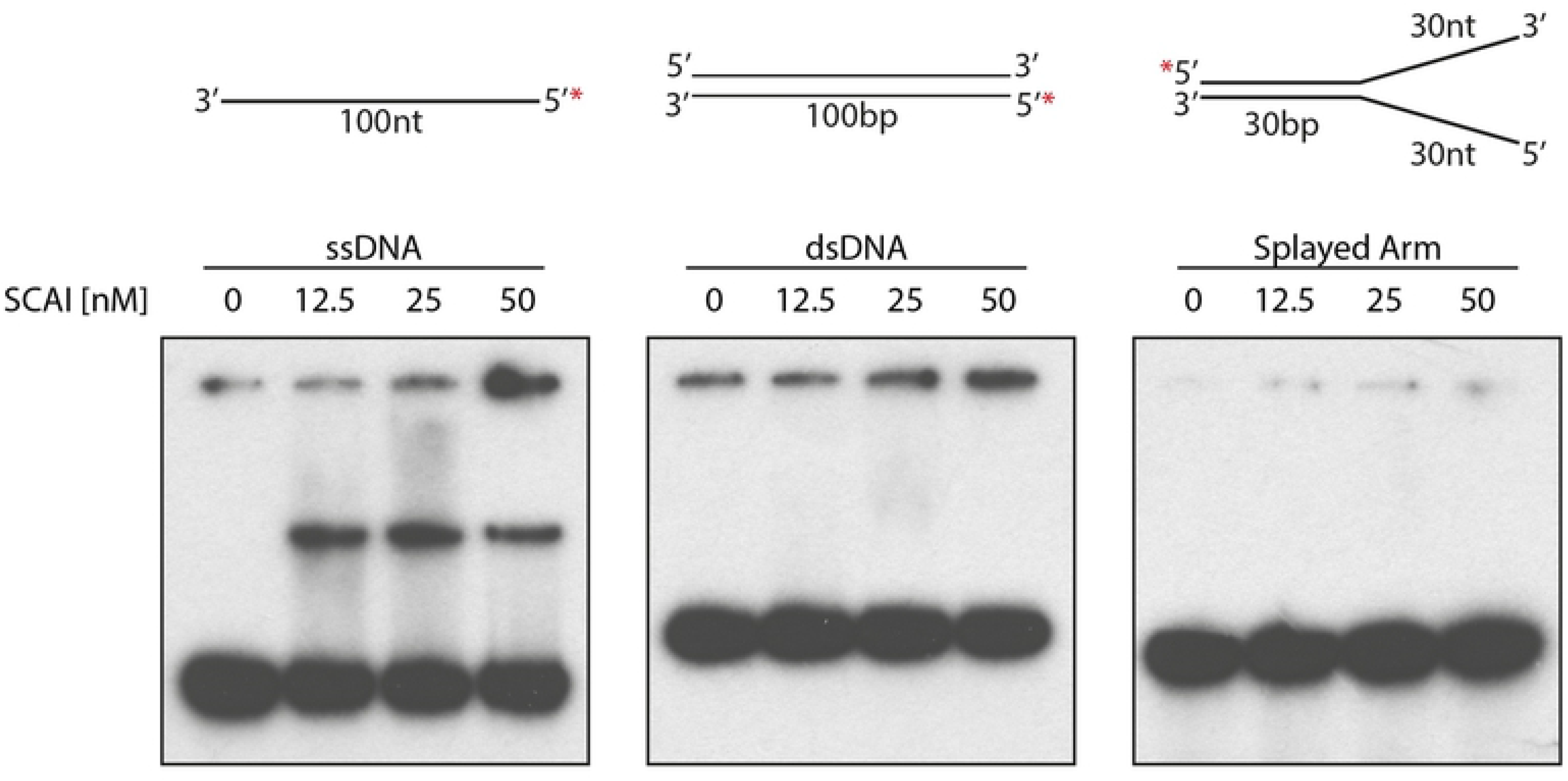
SCAI preferentially binds ssDNA over dsDNA. **A)** 5’-[^32^P]-labeled ssDNA, dsDNA or “splayed arm” DNA were incubated with purified recombinant SCAI at increasing concentrations and the reaction products were separated by acrylamide gel electrophoresis and visualized using autoradiography. Cartoons of the various substrates are shown on top of their respective gel. Representative results from 3 independents experiments.

## REFERENCES

Adeyemi RO, Willis NA, Elia AEH, Clairmont C, Li S, Wu X, D’Andrea AD, Scully R, Elledge SJ. 2021. The Protexin complex counters resection on stalled forks to promote homologous recombination and crosslink repair. Molecular Cell 0. doi:10.1016/j.molcel.2021.09.008

Auclair Y, Rouget R, Affar EB, Drobetsky EA. 2008. ATR kinase is required for global genomic nucleotide excision repair exclusively during S phase in human cells. Proc Natl Acad Sci USA 105:17896–17901. doi:10.1073/pnas.0801585105

Auclair Y, Rouget R, Belisle JM, Costantino S, Drobetsky EA. 2010. Requirement for functional DNA polymerase eta in genome-wide repair of UV-induced DNA damage during S phase. DNA Repair (Amst) 9:754–764. doi:10.1016/j.dnarep.2010.03.013

Bélanger F, Angers J-P, Fortier E, Hammond-Martel I, Costantino S, Drobetsky E, Wurtele H. 2015. Mutations in Replicative Stress Response Pathways Are Associated with S Phase-Specific Defects in Nucleotide Excision Repair. J Biol Chem. doi:10.1074/jbc.M115.685883

Bélanger F, Fortier E, Dubé M, Lemay J-F, Buisson R, Masson J-Y, Elsherbiny A, Costantino S, Carmona E, Mes-Masson A-M, Wurtele H, Drobetsky E. 2018. Replication Protein A Availability During DNA Replication Stress Is a Major Determinant of Cisplatin Resistance in Ovarian Cancer Cells. Cancer Res. doi:10.1158/0008-5472.CAN-18-0618

Brandt DT, Baarlink C, Kitzing TM, Kremmer E, Ivaska J, Nollau P, Grosse R. 2009. SCAI acts as a suppressor of cancer cell invasion through the transcriptional control of beta1-integrin. Nat Cell Biol 11:557–568. doi:10.1038/ncb1862

Branon TC, Bosch JA, Sanchez AD, Udeshi ND, Svinkina T, Carr SA, Feldman JL, Perrimon N, Ting AY. 2018. Efficient proximity labeling in living cells and organisms with TurboID. Nat Biotechnol 36:880–887. doi:10.1038/nbt.4201

Branzei D, Foiani M. 2009. The checkpoint response to replication stress. DNA Repair 8:1038– 1046. doi:16/j.dnarep.2009.04.014

Branzei D, Foiani M. 2007. Template Switching: From Replication Fork Repair to Genome Rearrangements. Cell 131:1228–1230. doi:16/j.cell.2007.12.007

Branzei D, Vanoli F, Foiani M. 2008. SUMOylation regulates Rad18-mediated template switch. Nature 456:915–920. doi:10.1038/nature07587

Buisson R, Lawrence MS, Benes CH, Zou L. 2017. APOBEC3A and 3B Activities Render Cancer Cells Susceptible to ATR Inhibition. Cancer Res 77:4567–4578. doi:10.1158/0008-5472.CAN-16-3389

Byun TS, Pacek M, Yee M, Walter JC, Cimprich KA. 2005. Functional uncoupling of MCM helicase and DNA polymerase activities activates the ATR-dependent checkpoint. Genes Dev 19:1040–1052. doi:10.1101/gad.1301205

Cantor SB. 2021. Revisiting the BRCA-pathway through the lens of replication gap suppression: “Gaps determine therapy response in BRCA mutant cancer.” DNA Repair 107:103209. doi:10.1016/j.dnarep.2021.103209

Chapman JR, Barral P, Vannier J-B, Borel V, Steger M, Tomas-Loba A, Sartori AA, Adams IR, Batista FD, Boulton SJ. 2013. RIF1 is essential for 53BP1-dependent nonhomologous end joining and suppression of DNA double-strand break resection. Mol Cell 49:858– 871. doi:10.1016/j.molcel.2013.01.002

Chen H, Lisby M, Symington LS. 2013. RPA coordinates DNA end resection and prevents formation of DNA hairpins. Mol Cell 50:10.1016/j.molcel.2013.04.032. doi:10.1016/j.molcel.2013.04.032

Chiang T-WW, le Sage C, Larrieu D, Demir M, Jackson SP. 2016. CRISPR-Cas9(D10A) nickase-based genotypic and phenotypic screening to enhance genome editing. Sci Rep 6:24356. doi:10.1038/srep24356

Cho KF, Branon TC, Udeshi ND, Myers SA, Carr SA, Ting AY. 2020. Proximity labeling in mammalian cells with TurboID and split-TurboID. Nat Protoc 15:3971–3999. doi:10.1038/s41596-020-0399-0

Choo JAMY, Schlösser D, Manzini V, Magerhans A, Dobbelstein M. 2020. The integrated stress response induces R-loops and hinders replication fork progression. Cell Death Dis 11. doi:10.1038/s41419-020-2727-2

Cong K, Peng M, Kousholt AN, Lee WTC, Lee S, Nayak S, Krais J, VanderVere-Carozza PS, Pawelczak KS, Calvo J, Panzarino NJ, Jonkers J, Johnson N, Turchi JJ, Rothenberg E, Cantor SB. 2021. Replication gaps are a key determinant of PARP inhibitor synthetic lethality with BRCA deficiency. Molecular Cell. doi:10.1016/j.molcel.2021.06.011

Costa RMA, Chiganças V, Galhardo R da S, Carvalho H, Menck CFM. 2003. The eukaryotic nucleotide excision repair pathway. Biochimie 85:1083–1099.

Diffley JFX. 2004. Regulation of early events in chromosome replication. Curr Biol 14:R778–786. doi:10.1016/j.cub.2004.09.019

Duan H, Mansour S, Reed R, Gillis MK, Parent B, Liu B, Sztupinszki Z, Birkbak N, Szallasi Z, Elia AEH, Garber JE, Pathania S. 2020. E3 ligase RFWD3 is a novel modulator of stalled fork stability in BRCA2-deficient cells. J Cell Biol 219. doi:10.1083/jcb.201908192

Dubois M-L, Bastin C, Lévesque D, Boisvert F-M. 2016. Comprehensive Characterization of Minichromosome Maintenance Complex (MCM) Protein Interactions Using Affinity and Proximity Purifications Coupled to Mass Spectrometry. J Proteome Res 15:2924–2934. doi:10.1021/acs.jproteome.5b01081

Elia AEH, Wang DC, Willis NA, Boardman AP, Hajdu I, Adeyemi RO, Lowry E, Gygi SP, Scully R, Elledge SJ. 2015. RFWD3-Dependent Ubiquitination of RPA Regulates Repair at Stalled Replication Forks. Mol Cell 60:280–293. doi:10.1016/j.molcel.2015.09.011

Forment JV, Jackson SP. 2015. A flow cytometry-based method to simplify the analysis and quantification of protein association to chromatin in mammalian cells. Nat Protoc 10:1297–1307. doi:10.1038/nprot.2015.066

Gallina I, Hendriks IA, Hoffmann S, Larsen NB, Johansen J, Colding-Christensen CS, Schubert L, Sellés-Baiget S, Fábián Z, Kühbacher U, Gao AO, Räschle M, Rasmussen S, Nielsen ML, Mailand N, Duxin JP. 2021. The ubiquitin ligase RFWD3 is required for translesion DNA synthesis. Mol Cell 81:442–458.e9. doi:10.1016/j.molcel.2020.11.029

Garzón J, Ursich S, Lopes M, Hiraga S-I, Donaldson AD. 2019. Human RIF1-Protein Phosphatase 1 Prevents Degradation and Breakage of Nascent DNA on Replication Stalling. Cell Rep 27:2558–2566.e4. doi:10.1016/j.celrep.2019.05.002

Giannattasio M, Follonier C, Tourrière H, Puddu F, Lazzaro F, Pasero P, Lopes M, Plevani P, Muzi-Falconi M. 2010. EXO1 competes with repair synthesis, converts NER intermediates to long ssDNA gaps, and promotes checkpoint activation. Mol Cell 40:50– 62. doi:10.1016/j.molcel.2010.09.004

Goodman MF, Woodgate R. 2013. Translesion DNA Polymerases. Cold Spring Harb Perspect Biol 5:a010363. doi:10.1101/cshperspect.a010363

Hansen RK, Mund A, Poulsen SL, Sandoval M, Klement K, Tsouroula K, Tollenaere MAX, Räschle M, Soria R, Offermanns S, Worzfeld T, Grosse R, Brandt DT, Rozell B, Mann M, Cole F, Soutoglou E, Goodarzi AA, Daniel JA, Mailand N, Bekker-Jensen S. 2016. SCAI promotes DNA double-strand break repair in distinct chromosomal contexts. Nat Cell Biol 18:1357–1366. doi:10.1038/ncb3436

He Z, Henricksen LA, Wold MS, Ingles CJ. 1995. RPA involvement in the damage-recognition and incision steps of nucleotide excision repair. Nature 374:566–569. doi:10.1038/374566a0

Hesketh GG, Youn J-Y, Samavarchi-Tehrani P, Raught B, Gingras A-C. 2017. Parallel Exploration of Interaction Space by BioID and Affinity Purification Coupled to Mass Spectrometry. Methods Mol Biol 1550:115–136. doi:10.1007/978-1-4939-6747-6_10

Hiraga S-I, Ly T, Garzón J, Hořejší Z, Ohkubo Y-N, Endo A, Obuse C, Boulton SJ, Lamond AI, Donaldson AD. 2017. Human RIF1 and protein phosphatase 1 stimulate DNA replication origin licensing but suppress origin activation. EMBO Rep 18:403–419. doi:10.15252/embr.201641983

Hristova RH, Stoynov SS, Tsaneva IR, Gospodinov AG. 2020. Deregulated levels of RUVBL1 induce transcription-dependent replication stress. Int J Biochem Cell Biol 128:105839. doi:10.1016/j.biocel.2020.105839

Isobe S-Y, Nagao K, Nozaki N, Kimura H, Obuse C. 2017. Inhibition of RIF1 by SCAI Allows BRCA1-Mediated Repair. Cell Rep 20:297–307. doi:10.1016/j.celrep.2017.06.056

Iyer DR, Rhind N. 2017. The Intra-S Checkpoint Responses to DNA Damage. Genes (Basel) 8. doi:10.3390/genes8020074

Joung J, Konermann S, Gootenberg JS, Abudayyeh OO, Platt RJ, Brigham MD, Sanjana NE, Zhang F. 2017. Genome-scale CRISPR-Cas9 knockout and transcriptional activation screening. Nat Protoc 12:828–863. doi:10.1038/nprot.2017.016

Kechin A, Boyarskikh U, Kel A, Filipenko M. 2017. cutPrimers: A New Tool for Accurate Cutting of Primers from Reads of Targeted Next Generation Sequencing. J Comput Biol 24:1138–1143. doi:10.1089/cmb.2017.0096

Kim JJ, Lee SY, Choi J-H, Woo HG, Xhemalce B, Miller KM. 2020. PCAF-Mediated Histone Acetylation Promotes Replication Fork Degradation by MRE11 and EXO1 in BRCA-Deficient Cells. Molecular Cell 80:327–344.e8. doi:10.1016/j.molcel.2020.08.018

Knowlton JJ, Gestaut D, Ma B, Taylor G, Seven AB, Leitner A, Wilson GJ, Shanker S, Yates NA, Prasad BVV, Aebersold R, Chiu W, Frydman J, Dermody TS. 2021. Structural and functional dissection of reovirus capsid folding and assembly by the prefoldin-TRiC/CCT chaperone network. Proc Natl Acad Sci U S A 118:e2018127118. doi:10.1073/pnas.2018127118

Kolinjivadi AM, Sannino V, Antoni AD, Zadorozhny K, Kilkenny M, Técher H, Baldi G, Shen R, Ciccia A, Pellegrini L, Krejci L, Costanzo V. 2017a. Smarcal1-Mediated Fork Reversal Triggers MRE11-Dependent Degradation of Nascent DNA in the Absence of Brca2 and Stable Rad51 Nucleofilaments. Molecular Cell 67:867–881.e7. doi:10.1016/j.molcel.2017.07.001

Kolinjivadi AM, Sannino V, de Antoni A, Técher H, Baldi G, Costanzo V. 2017b. Moonlighting at replication forks -a new life for homologous recombination proteins BRCA1, BRCA2 and RAD51. FEBS Lett 591:1083–1100. doi:10.1002/1873-3468.12556

Lambert É, Babeu J-P, Simoneau J, Raisch J, Lavergne L, Lévesque D, Jolibois É, Avino M, Scott MS, Boudreau F, Boisvert F-M. 2020. Human Hepatocyte Nuclear Factor 4-α Encodes Isoforms with Distinct Transcriptional Functions. Mol Cell Proteomics 19:808– 827. doi:10.1074/mcp.RA119.001909

Lemaçon D, Jackson J, Quinet A, Brickner JR, Li S, Yazinski S, You Z, Ira G, Zou L, Mosammaparast N, Vindigni A. 2017. MRE11 and EXO1 nucleases degrade reversed forks and elicit MUS81-dependent fork rescue in BRCA2-deficient cells. Nat Commun 8:860. doi:10.1038/s41467-017-01180-5

Li W, Xu H, Xiao T, Cong L, Love MI, Zhang F, Irizarry RA, Liu JS, Brown M, Liu XS. 2014. MAGeCK enables robust identification of essential genes from genome-scale CRISPR/Cas9 knockout screens. Genome Biol 15:554. doi:10.1186/s13059-014-0554-4

Lindahl T. 1993. Instability and decay of the primary structure of DNA. Nature 362:709–715. doi:10.1038/362709a0

Llamas E, Torres-Montilla S, Lee HJ, Barja MV, Schlimgen E, Dunken N, Wagle P, Werr W, Zuccaro A, Rodríguez-Concepción M, Vilchez D. 2021. The intrinsic chaperone network of Arabidopsis stem cells confers protection against proteotoxic stress. Aging Cell 20:e13446. doi:10.1111/acel.13446

Maréchal A, Zou L. 2015. RPA-coated single-stranded DNA as a platform for post-translational modifications in the DNA damage response. Cell Res 25:9–23. doi:10.1038/cr.2014.147

Martín-Cófreces NB, Valpuesta JM, Sánchez-Madrid F. 2021. Folding for the Immune Synapse: CCT Chaperonin and the Cytoskeleton. Front Cell Dev Biol 9:658460. doi:10.3389/fcell.2021.658460

Mattarocci S, Shyian M, Lemmens L, Damay P, Altintas DM, Shi T, Bartholomew CR, Thomä NH, Hardy CFJ, Shore D. 2014. Rif1 controls DNA replication timing in yeast through the PP1 phosphatase Glc7. Cell Rep 7:62–69. doi:10.1016/j.celrep.2014.03.010

Mejlvang J, Feng Y, Alabert C, Neelsen KJ, Jasencakova Z, Zhao X, Lees M, Sandelin A, Pasero P, Lopes M, Groth A. 2014. New histone supply regulates replication fork speed and PCNA unloading. J Cell Biol 204:29–43. doi:10.1083/jcb.201305017

Mellacheruvu D, Wright Z, Couzens AL, Lambert J-P, St-Denis NA, Li T, Miteva YV, Hauri S, Sardiu ME, Low TY, Halim VA, Bagshaw RD, Hubner NC, Al-Hakim A, Bouchard A, Faubert D, Fermin D, Dunham WH, Goudreault M, Lin Z-Y, Badillo BG, Pawson T, Durocher D, Coulombe B, Aebersold R, Superti-Furga G, Colinge J, Heck AJR, Choi H, Gstaiger M, Mohammed S, Cristea IM, Bennett KL, Washburn MP, Raught B, Ewing RM, Gingras A-C, Nesvizhskii AI. 2013. The CRAPome: a contaminant repository for affinity purification-mass spectrometry data. Nat Methods 10:730–736. doi:10.1038/nmeth.2557

Mi H, Muruganujan A, Huang X, Ebert D, Mills C, Guo X, Thomas PD. 2019. Protocol Update for large-scale genome and gene function analysis with the PANTHER classification system (v.14.0). Nat Protoc 14:703–721. doi:10.1038/s41596-019-0128-8

Mijic S, Zellweger R, Chappidi N, Berti M, Jacobs K, Mutreja K, Ursich S, Ray Chaudhuri A, Nussenzweig A, Janscak P, Lopes M. 2017. Replication fork reversal triggers fork degradation in BRCA2-defective cells. Nat Commun 8. doi:10.1038/s41467-017-01164-5

Nakazawa Y, Yamashita S, Lehmann AR, Ogi T. 2010. A semi-automated non-radioactive system for measuring recovery of RNA synthesis and unscheduled DNA synthesis using ethynyluracil derivatives. DNA Repair (Amst) 9:506–516. doi:10.1016/j.dnarep.2010.01.015

Nayak S, Calvo JA, Cong K, Peng M, Berthiaume E, Jackson J, Zaino AM, Vindigni A, Hadden MK, Cantor SB. 2020. Inhibition of the translesion synthesis polymerase REV1 exploits replication gaps as a cancer vulnerability. Sci Adv 6:eaaz7808. doi:10.1126/sciadv.aaz7808

Neelsen KJ, Lopes M. 2015. Replication fork reversal in eukaryotes: from dead end to dynamic response. Nat Rev Mol Cell Biol 16:207–220. doi:10.1038/nrm3935

Oakley GG, Patrick SM. 2010. Replication protein A: directing traffic at the intersection of replication and repair. Front Biosci (Landmark Ed) 15:883–900.

Panzarino NJ, Krais JJ, Cong K, Peng M, Mosqueda M, Nayak SU, Bond SM, Calvo JA, Doshi MB, Bere M, Ou J, Deng B, Zhu LJ, Johnson N, Cantor SB. 2021. Replication Gaps Underlie BRCA Deficiency and Therapy Response. Cancer Res 81:1388–1397. doi:10.1158/0008-5472.CAN-20-1602

Parrilla-Castellar ER, Arlander SJH, Karnitz L. 2004. Dial 9-1-1 for DNA damage: the Rad9-Hus1-Rad1 (9-1-1) clamp complex. DNA Repair (Amst) 3:1009–1014. doi:10.1016/j.dnarep.2004.03.032

Piberger AL, Bowry A, Kelly RDW, Walker AK, González-Acosta D, Bailey LJ, Doherty AJ, Méndez J, Morris JR, Bryant HE, Petermann E. 2020. PrimPol-dependent single-stranded gap formation mediates homologous recombination at bulky DNA adducts. Nat Commun 11:5863. doi:10.1038/s41467-020-19570-7

Provencher DM, Lounis H, Champoux L, Tétrault M, Manderson EN, Wang JC, Eydoux P, Savoie R, Tonin PN, Mes-Masson AM. 2000. Characterization of four novel epithelial ovarian cancer cell lines. In Vitro Cell Dev Biol Anim 36:357–361. doi:10.1290/1071-2690(2000)036<0357:COFNEO>2.0.CO;2

Quinet A, Carvajal-Maldonado D, Lemacon D, Vindigni A. 2017. DNA Fiber Analysis: Mind the Gap! Meth Enzymol 591:55–82. doi:10.1016/bs.mie.2017.03.019

Quinet A, Tirman S, Cybulla E, Meroni A, Vindigni A. 2021. To skip or not to skip: choosing repriming to tolerate DNA damage. Molecular Cell 81:649–658. doi:10.1016/j.molcel.2021.01.012

Roux KJ, Kim DI, Raida M, Burke B. 2012. A promiscuous biotin ligase fusion protein identifies proximal and interacting proteins in mammalian cells. J Cell Biol 196:801–810. doi:10.1083/jcb.201112098

Sanjana NE, Shalem O, Zhang F. 2014. Improved vectors and genome-wide libraries for CRISPR screening. Nat Meth 11:783–784. doi:10.1038/nmeth.3047

Santocanale C, Diffley JF. 1998. A Mec1- and Rad53-dependent checkpoint controls late-firing origins of DNA replication. Nature 395:615–618. doi:10.1038/27001

Schlacher K, Christ N, Siaud N, Egashira A, Wu H, Jasin M. 2011. Double-Strand Break Repair Independent Role For BRCA2 In Blocking Stalled Replication Fork Degradation By MRE11. Cell 145:529–542. doi:10.1016/j.cell.2011.03.041

Segurado M, Diffley JFX. 2008. Separate roles for the DNA damage checkpoint protein kinases in stabilizing DNA replication forks. Genes Dev 22:1816–1827. doi:10.1101/gad.477208

Shaheen N, Akhtar J, Umer Z, Khan MHF, Bakhtiari MH, Saleem M, Faisal A, Tariq M. 2021. Polycomb Requires Chaperonin Containing TCP-1 Subunit 7 for Maintaining Gene Silencing in Drosophila. Front Cell Dev Biol 9:727972. doi:10.3389/fcell.2021.727972

Shalem O, Sanjana NE, Hartenian E, Shi X, Scott DA, Mikkelsen TS, Heckl D, Ebert BL, Root DE, Doench JG, Zhang F. 2014. Genome-scale CRISPR-Cas9 knockout screening in human cells. Science 343:84–87. doi:10.1126/science.1247005

Shanbhag NM, Rafalska-Metcalf IU, Balane-Bolivar C, Janicki SM, Greenberg RA. 2010. ATM-dependent chromatin changes silence transcription in cis to DNA double-strand breaks. Cell 141:970–981. doi:10.1016/j.cell.2010.04.038

Sogo JM, Lopes M, Foiani M. 2002. Fork reversal and ssDNA accumulation at stalled replication forks owing to checkpoint defects. Science 297:599–602. doi:10.1126/science.1074023

Tkáč J, Xu G, Adhikary H, Young JTF, Gallo D, Escribano-Díaz C, Krietsch J, Orthwein A, Munro M, Sol W, Al-Hakim A, Lin Z-Y, Jonkers J, Borst P, Brown GW, Gingras A-C, Rottenberg S, Masson J-Y, Durocher D. 2016. HELB Is a Feedback Inhibitor of DNA End Resection. Molecular Cell 61:405–418. doi:10.1016/j.molcel.2015.12.013

Toledo L, Neelsen KJ, Lukas J. 2017. Replication Catastrophe: When a Checkpoint Fails because of Exhaustion. Molecular Cell 66:735–749. doi:10.1016/j.molcel.2017.05.001

Toledo LI, Altmeyer M, Rask M-B, Lukas C, Larsen DH, Povlsen LK, Bekker-Jensen S, Mailand N, Bartek J, Lukas J. 2013. ATR prohibits replication catastrophe by preventing global exhaustion of RPA. Cell 155:1088–1103. doi:10.1016/j.cell.2013.10.043

Tsaalbi-Shtylik A, Moser J, Mullenders LHF, Jansen JG, de Wind N. 2014. Persistently stalled replication forks inhibit nucleotide excision repair in trans by sequestering Replication protein A. Nucleic Acids Res 42:4406–4413. doi:10.1093/nar/gkt1412

van den Heuvel D, Spruijt CG, González-Prieto R, Kragten A, Paulsen MT, Zhou D, Wu H, Apelt K, van der Weegen Y, Yang K, Dijk M, Daxinger L, Marteijn JA, Vertegaal ACO, Ljungman M, Vermeulen M, Luijsterburg MS. 2021. A CSB-PAF1C axis restores processive transcription elongation after DNA damage repair. Nat Commun 12:1342. doi:10.1038/s41467-021-21520-w

Wang B, Wang M, Zhang W, Xiao T, Chen C-H, Wu A, Wu F, Traugh N, Wang X, Li Z, Mei S, Cui Y, Shi S, Lipp JJ, Hinterndorfer M, Zuber J, Brown M, Li W, Liu XS. 2019. Integrative analysis of pooled CRISPR genetic screens using MAGeCKFlute. Nat Protoc 14:756–780. doi:10.1038/s41596-018-0113-7

Weng H, Feng X, Lan Y, Zheng Z. 2021. TCP1 regulates PI3K/AKT/mTOR signaling pathway to promote proliferation of ovarian cancer cells. J Ovarian Res 14:82. doi:10.1186/s13048-021-00832-x

Yam AY, Xia Y, Lin H-TJ, Burlingame A, Gerstein M, Frydman J. 2008. Defining the TRiC/CCT interactome links chaperonin function to stabilization of newly made proteins with complex topologies. Nat Struct Mol Biol 15:1255–1262. doi:10.1038/nsmb.1515

Yau EH, Rana TM. 2018. Next-Generation Sequencing of Genome-Wide CRISPR Screens. Methods Mol Biol 1712:203–216. doi:10.1007/978-1-4939-7514-3_13

Yekezare M, Gómez-González B, Diffley JFX. 2013. Controlling DNA replication origins in response to DNA damage – inhibit globally, activate locally. J Cell Sci 126:1297–1306. doi:10.1242/jcs.096701

Zellweger R, Dalcher D, Mutreja K, Berti M, Schmid JA, Herrador R, Vindigni A, Lopes M. 2015. Rad51-mediated replication fork reversal is a global response to genotoxic treatments in human cells. J Cell Biol 208:563–579. doi:10.1083/jcb.201406099

Zeman MK, Cimprich KA. 2014. Causes and consequences of replication stress. Nat Cell Biol 16:2–9. doi:10.1038/ncb2897

Zimmermann M, de Lange T. 2014. 53BP1: pro choice in DNA repair. Trends Cell Biol 24:108– 117. doi:10.1016/j.tcb.2013.09.003

